# Visualizing looping of two endogenous genomic loci using synthetic zinc-finger proteins with anti-FLAG and anti-HA frankenbodies in living cells

**DOI:** 10.1101/2021.06.16.448697

**Authors:** Yang Liu, Ning Zhao, Masato T. Kanemaki, Yotaro Yamamoto, Yoshifusa Sadamura, Yuma Ito, Makio Tokunaga, Timothy J. Stasevich, Hiroshi Kimura

## Abstract

In eukaryotic nuclei, chromatin loops mediated through cohesin are critical structures that regulate gene expression and DNA replication. Here we demonstrate a new method to visualize endogenous genomic loci using synthetic zinc-finger proteins harboring repeat epitope tags (ZF probes) for signal amplification via binding of tag-specific intracellular antibodies, or frankenbodies, fused with fluorescent proteins. We achieve this in two steps. First, we develop an anti-FLAG frankenbody that can bind FLAG-tagged proteins in diverse live-cell environments. The anti-FLAG frankenbody complements the anti-HA frankenbody, enabling two-color signal amplification from FLAG and HA-tagged proteins. Second, we develop a pair of cell-permeable ZF probes that specifically bind two endogenous chromatin loci predicted to be involved in chromatin looping. By coupling our anti-FLAG and anti-HA frankenbodies with FLAG- and HA-tagged ZF probes, we simultaneously visualize the dynamics of the two loci in single living cells. This reveals close association between the two loci in the majority of cells, but the loci markedly separate upon the triggered degradation of the cohesin subunit RAD21. Our ability to image two endogenous genomic loci simultaneously in single living cells provides a proof-of-principle that ZF probes coupled with frankenbodies are useful new tools for exploring genome dynamics in multiple colors.

## INTRODUCTION

During cell cycle progression and cell differentiation, chromosomes undergo dynamic changes in their structure and organization. In the interphase nucleus, for example, chromatin loops are critical structures that regulate gene expression and DNA replication (Baumann, 2020; Hansen et al., 2017). Chromosome conformation capture (3C; (Dekker et al., 2002)) and Hi-C (Lieberman-Aiden et al., 2009) analyses have revealed chromatin loops help separate active chromatin domains from inactive ones (Bickmore, 2012). The formation of chromatin loops creates chromosome clusters, also known as chromosome compartments, that span several hundred Kb to a few Mb (Dixon et al., 2012; Nora et al., 2012), contain topologically associating domains (TADs), CTCF (CCCTC-binding factor) binding sites, and cohesin complexes (Pombo & Dillon, 2015).

So far, chromatin looping has mainly been investigated in fixed cells using Hi-C (Dixon et al., 2012; Nora et al., 2012) and fluorescence in situ hybridization (FISH; (Dillon et al., 1997; Gizzi et al., 2019; Joyce et al., 2012; Wijgerde et al., 1995; Woglar et al., 2020)). While powerful, these assays require fixation, leaving the dynamics of chromatin looping unclear. To better resolve these dynamics, including changes in promoter-enhancer contacts in association with gene expression and formation of aberrant contacts that can cause diseases, a sequence-specific live-cell imaging technology would be useful (Shaban et al., 2020).

To image genomic loci in living cells, several methods have been developed. The pioneering work labeled specific or non-specific genomic loci by inserting tens or hundreds of lac or tet operators that could be bound by fluorescent Lac or Tet repressors. This created ultra-bright fluorescent spots that marked the location of the repeats within the genome so they could be tracked for long periods of time using standard fluorescence microscopy (Michaelis et al., 1997; Robinett et al., 1996). Such artificial repeat-based methods can yield high contrast images because hundreds of fluorophores can be present in a single spot. In this approach, however, extensive genomic manipulation is essential and the insertion may affect gene regulation around the inserted locus. Recently, endogenous loci have been visualized using zinc-finger (ZF) proteins, transcription activator-like effectors (TALEs), and CRISPR/nuclease-deficient Cas9 (dCas9) systems, without disturbing cell growth and embryo development (Anton et al., 2014; Lindhout et al., 2007; Ma et al., 2018; Miyanari et al., 2013). Whereas most probes target repeat sequences, specific genomic loci can also be labeled with multiple unique probes (Chen et al., 2013). CRISPR/dCas9 has been a convenient and popular method because probe specificity is easily modulated by altering sgRNA sequence (Ran et al., 2013), whereas it is essential to design and construct zinc-finger proteins and TALEs for each specific sequence. However, the strong DNA-binding nature of some probes, including Lac repressor and dCas9 with sgRNA, could interfere with DNA replication and transcription (Doi et al., 2021; Garcia-Bloj et al., 2016; Jiang et al., 2015; Qi et al., 2013; Rinaldi et al., 2017), depending on their target location. By contrast, ParB and ZF proteins without extra functional group fusions have been shown to interfere less with such events (Garriga-Canut et al., 2012; Gersbach et al., 2014; Imanishi et al., 2000; Isalan et al., 1997; Saad et al., 2014). The low toxicity of ZF probes has made ZF-based therapies popular in ongoing human clinical trials (Lee et al., 2016; Perez et al., 2008; Tebas et al., 2014; Wilen et al., 2011).

In this study, we develop technology for genome visualization based on ZF DNA-binding proteins harboring repeat epitope tags for signal amplification by epitope-specific intracellular antibodies. To achieve this, we first developed an anti-FLAG “frankenbody” – a chimeric single chain variable fragment that binds FLAG epitopes in living cells – to complement our recently developed anti-HA frankenbody (Zhao et al., 2019). Together, these two frankenbodies can bind and label HA- and FLAG-tagged proteins with two distinct fluorophores in single living cells. We next generated a pair of cell-permeable ZF DNA-binding probes (Barrow et al., 2012; Choo et al., 1994; Gaj & Liu, 2015; Gaj et al., 2014; Mino et al., 2008; Sera, 2010) and tagged them with 10× HA or FLAG epitopes. When these primary ZF probes are delivered into living cells that express anti-HA and anti-FLAG frankenbodies with different fluorescent proteins as secondary probes, two distinct endogenous genomic loci can be simultaneously marked and tracked. Using this system, we followed two genomic regions that are near cohesin binding sites and show close contact in cellular 3D space according to HiC. Upon triggered cohesin degradation (Natsume et al., 2016), the two regions became more separated, confirming that the strategy can be used to monitor cohesin-mediated chromatin contacts. We anticipate our ZF-based technology can now be extended to monitor other endogenous genomic loci and investigate the dynamics of higher-order chromatin structures in a multiplexed manner.

## Materials and methods

### Construction of plasmids for anti-FLAG frankenbody development and characterization

Variable regions of antibody heavy and light chains encoding an anti-FLAG antibody were amplified by PCR and the nucleotide sequences were determined (Sato et al., 2013). Wildtype and chimeric anti-FLAG scFv genes were synthesized as gblocks. The scFv gblocks were ligated with linearized anti-HA frankenbody-mEGFP (Addgene # 129590) by EcoRI through Gibson assembly (anti-FLAG FB-mEGFP). Anti-FLAG frankenbody fused with mRuby2 was constructed by Gibson assembly by ligating NotI-linearized anti-FLAG FB-mEGFP with mRuby2 amplified from a previously built plasmid (4×HA-mRuby-Kv2.1). Anti-FLAG frankenbody fused with SNAP-tag and HaloTag plasmids were constructed by ligating previously built plasmids, anti-HA FB-SNAP and anti-HA FB-Halo (Addgene #129592), cut by EcoRI and combined with anti-FLAG frankenbody gblocks by Gibson assembly. Anti-HA frankenbody mCherry and anti-FLAG iRFP were constructed by inserting PCR-amplified mCherry and iRPF sequences harboring SalI and BsrGI sites into a fragment derived from anti-FLAG FB-EGFP digested with the same enzymes by ligation. Previously built plasmids derived from pmCherry-N1 (Takara-Clontech) and piRFP (a gift from Michael Davidson & Vladislav Verkhusha; Addgene plasmid # 31857 (Filonov et al., 2011)) were used as the template of PCR. These anti-FLAG frankenbody plasmid constructs will be available at Addgene.

4×FLAG-mCh-H2B and 4×FLAG-mCh-β-actin plasmids were built by Gibson assembly by ligating a synthesized 4×FLAG gblock with previously built 1×HA-mCh-H2B or 1×HA-mCh-β-actin plasmids cut by AgeI and NotI. 4×FLAG-sfGFP-H2B was built by ligating 4×FLAG-mCh-H2B cut by NotI and BglII with sfGFP-H2B amplified from an Addgene plasmid, sfGFP-H2B-C-10 (#56367). Mito-mCh-1×FLAG and Mito-mCh-smFLAG were constructed by ligating 1×FLAG synthesized by overlapping PCR and smFLAG amplified from smFLAG-KDM5B-24×MS2 (Addgene # 81084) with previously built Mito-mCh-1×HA cut by BglII and BamHI through Gibson Assembly.

The gblocks were synthesized by Integrated DNA Technologies and the recombinant plasmids were sequence verified by Quintara Biosciences. All plasmids used for imaging were prepared using a NucleoBond Xtra Midi EF kit (Macherey-Nagel) with a final concentration of about 1 mg/mL.

### Cell culture and transfection

U2OS cells (ATCC HTB-96) were cultured in DMEM medium (Thermo Fisher Scientific) supplemented with 10% (v/v) fetal bovine serum (FBS; Altas Biologicals), 1 mM L-glutamine and 1% (v/v) penicillin-streptomycin (Thermo Fisher Scientific) at 37°C, in a cell culture incubator under 5% of CO_2_. HCT116-RAD21-mAID-mClover cells (Natsume et al., 2016) were cultured in McCoy’s 5A (Modified) Medium with 10% FBS (Thermo Fisher Scientific) and 1% Glutamine-Penicillin-Streptomycin solution (Sigma-Aldrich) at 37°C in a cell culture incubator under 5% of CO_2_. HCT116 (ATCC), HEK-293T (obtained from Kei Fujinaga at Sapporo Medical School in 1980s) and A9 cells (obtained from Nobuo Takagi at Hokkaido University in 1980s) were cultured in DMEM (Nacalai Tesque) with 10% FCS (Thermo Fisher Scientific) and 1% GPS solution (Sigma-Aldrich) at 37°C in a cell culture incubator under 5% of CO_2_. Lipofectamine LTX reagent with PLUS reagent (Thermo Fisher Scientific) or Lipofectamine 3000 (Thermo Fisher Scientific) was used for transfection following the manufacturer’s instructions.

### Preparing cells for imaging single-mRNA translation dynamics

Imaging reagents (smFLAG-KDM5B-24×MS2/anti-FLAG FB-GFP and purified MCP-HaloTag protein) needed for nascent chain tracking and puromycin treatment were bead loaded into cells as previously described (Zhao et al., 2019). Briefly, U2OS cells were plated on 35 mm MatTek chambers (MatTek) the day before imaging. On the imaging day, the medium in the MatTek chambers was changed to Opti-MEM (Thermo Fisher Scientific) with 10% FBS. The cells were incubated in the Opti-MEM for 20 min. Then 4 µL of a mixture of plasmids (1 µg of smFLAG-KDM5B-24×MS2 and 0.5 µg of anti-FLAG FB-GFP) and purified MCP-HaloTag (130 ng) in PBS was pipetted on top of the cells after removing the Opti-medium from the MatTek chamber, and ~106 µm glass beads (Sigma Aldrich) were evenly distributed on top. The chamber was then tapped firmly seven times, and Opti-medium was added back to the cells (Cialek et al., 2021). 3 h post bead loading, the cells were stained in 1 mL of 0.2 µM of JF646-HaloTag ligand (Grimm et al., 2015) diluted in phenol-red-free complete DMEM. After 20 min of staining, the cells were washed three times in phenol-red-free complete DMEM to remove glass beads and excess dyes. The cells were then ready for imaging.

### Fluorescence recovery after photobleaching experiments

Fluorescence recovery after photobleaching (FRAP) assays were performed on cells transiently transfected with 4×FLAG-mCh-H2B and anti-FLAG FB-mEGFP 18~22 h before FRAP. The images were acquired using an Olympus IX81 spinning disk confocal (CSU22 head) microscope coupled to a Phasor photomanipulation unit (Intelligent Imaging Innovations) with a 100×oil immersion objective (NA 1.40). Before photobleaching, 20 frames were acquired with a 1s time interval. The images were captured using a 488 nm laser (0.77 mW) followed by a 561 nm (0.42 mW) laser. The laser exposure time was adjusted according to the fluorescence intensity. The spinning disk was set up at a 1 × 1 spin rate. Images were acquired with a Photometrics Cascade II electron multiplying-coupled charge device (EM-CCD) using SlideBook software (Intelligent Imaging Innovations). After acquiring pre-FRAP images, the 488 nm laser (from the Phasor unit for photobleaching) was set to 17 mW with a 100 ms exposure time to photobleach a circular region in the nucleus. After photobleaching, 30 images were captured without delay, and then an additional 100 images were acquired with a 5s delay using the same imaging settings as the pre-FRAP images. The fluorescence intensity through time of the photobleached spot was exported using Slidebook software (Intelligent Imaging Innovations). The fluorescence intensity of the nucleus and background were obtained by ImageJ. The FRAP curve and *t_half_* were obtained using easyFRAP-web (Koulouras et al., 2018), according to the website instructions.

### Imaging conditions for FLAG colocalization experiments and nascent chain tracking

For the co-localization assays in Figs. 1B, 1C and 2A, images were acquired using an Olympus IX81 spinning disk confocal (CSU22 head) microscope using a 100× oil immersion objective (NA 1.40) under the following conditions: 488 nm (0.77 mW) and 561 nm (0.42 mW) sequential imaging for 50 time points without delay at a single plane for Figs. 1B and 1C, and for five time points without delay with 13 z-slices to cover the whole cell body for each time point for Fig. 2A; 1 × 1 spin rate, exposure time adjusted by cell brightness. Images were acquired with a Photometrics Cascade II EM-CCD camera using SlideBook software (Intelligent Imaging Innovations). The displayed images in Figs. 1B and 1C were generated by averaging 50 time points and the images in Fig. 2A were generated by averaging five time points and then max-projecting all z-slices in ImageJ.

**Fig. 1.**
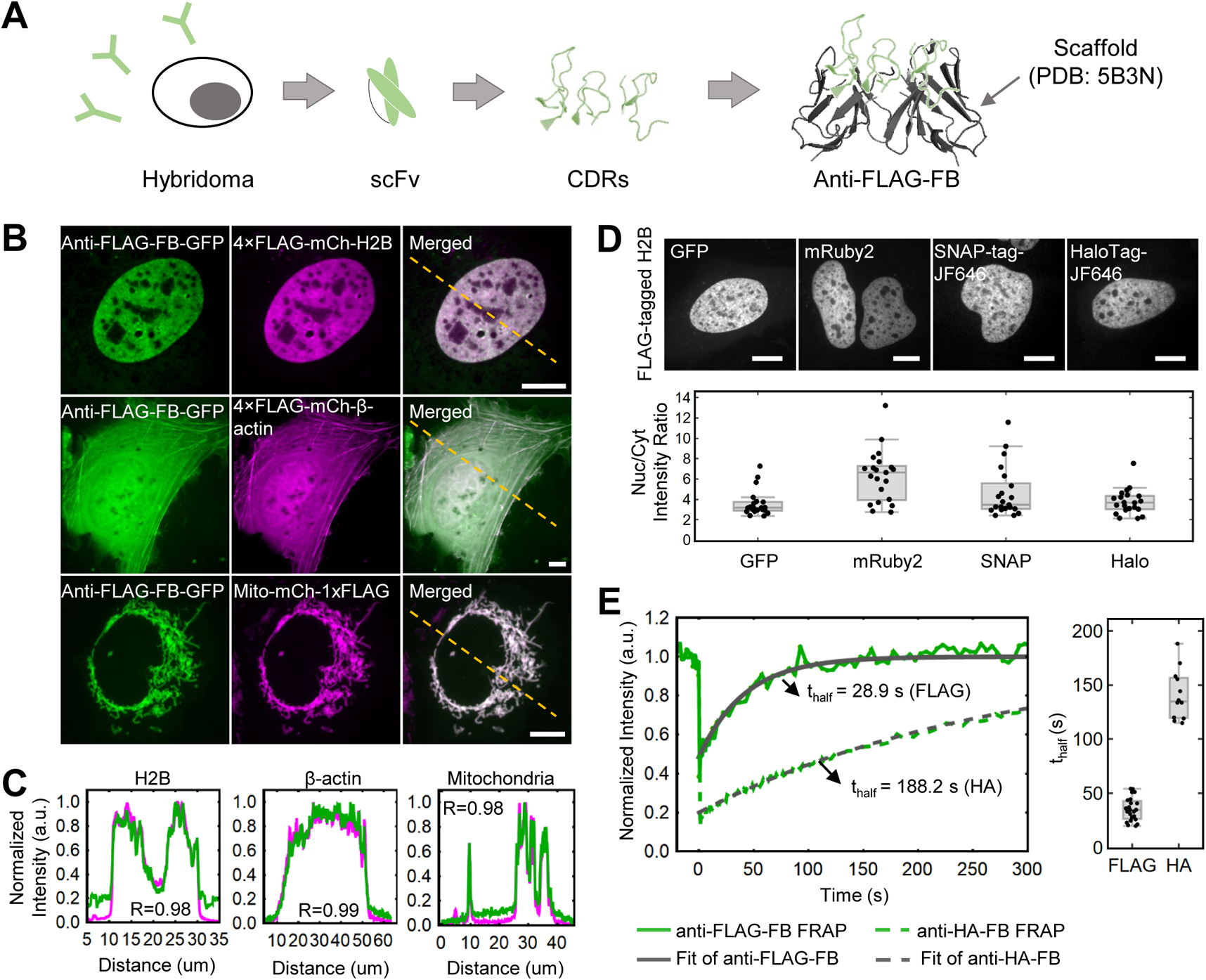
Development of an anti-FLAG frankenbody to label FLAG-tagged proteins in living cells. (**A**) A schematic showing how to design anti-FLAG frankenbodies (anti-FLAG-FB) using anti-FLAG-scFv CDRs and stable scFv scaffolds. (**B**) Representative cells showing the respective localization of the anti-FLAG frankenbody in living U2OS cells co-expressing FLAG-tagged proteins of interest (POIs), including nuclear protein histone H2B (n=20 cells), cytoplasmic protein β-actin (n=9 cells) and mitochondria (n=30 cells), with the FLAG tag located at either the N- or C-terminus (anti-FLAG frankenbody, green; FLAG-tagged mCh-POIs, magenta). (**C**) Normalized fluorescence intensity plots of the yellow dashed lines shown in (**B**) for both frankenbody (green) and FLAG-tagged mCh-POI channels (magenta). The intensity was normalized by setting the minimum and maximum intensities to 0 and 1, respectively. (**D**) Upper panel: Representative cell images of anti-FLAG frankenbody fused to multiple fluorescent fusion proteins (GFP, mRuby2, SNAP-tag/JF646 and HaloTag/JF646) and specifically labeleling FLAG-tagged nuclear protein H2B (FLAG-tagged H2B). Lower panel: Box plot showing the nuclear to cytoplasmic fluorescence intensity ratio (Nuc/Cyt) for all cells imaged in upper panel; 21 cells in one independent experiment for each fluorophore. (**E**) Left: Plot of the anti-FLAG and anti-HA frankenbody FRAP data with fitted curves for representative cells (the green solid line is the FRAP recovery curve for the anti-FLAG frankenbody and the gray solid line is the fitted curve; the green dashed line is the FRAP recovery curve for the anti-HA frankenbody and gray dashed line is the fitted curve); Right: Box plot showing the recovery halftimes of all FRAP experiments for the anti-FLAG frankenbody (n=23 cells in 2 independent experiments) and the anti-HA frankenbody. Fits from 23 cells reveal the mean FRAP recovery halftime (*t_half_*) of the anti-FLAG frankenbody is 35.3±2.2 s (cell-to-cell SEM), while that of the anti-HA frankenbody is 141±7 s. Scale bars: 10 µm. For box plots (**A** and **E**), the center lines show the medians; the boxes indicate 25-75%; whiskers extend 1.5 times the interquartile range from the 25th and 75th percentiles.

**Fig. 2.**
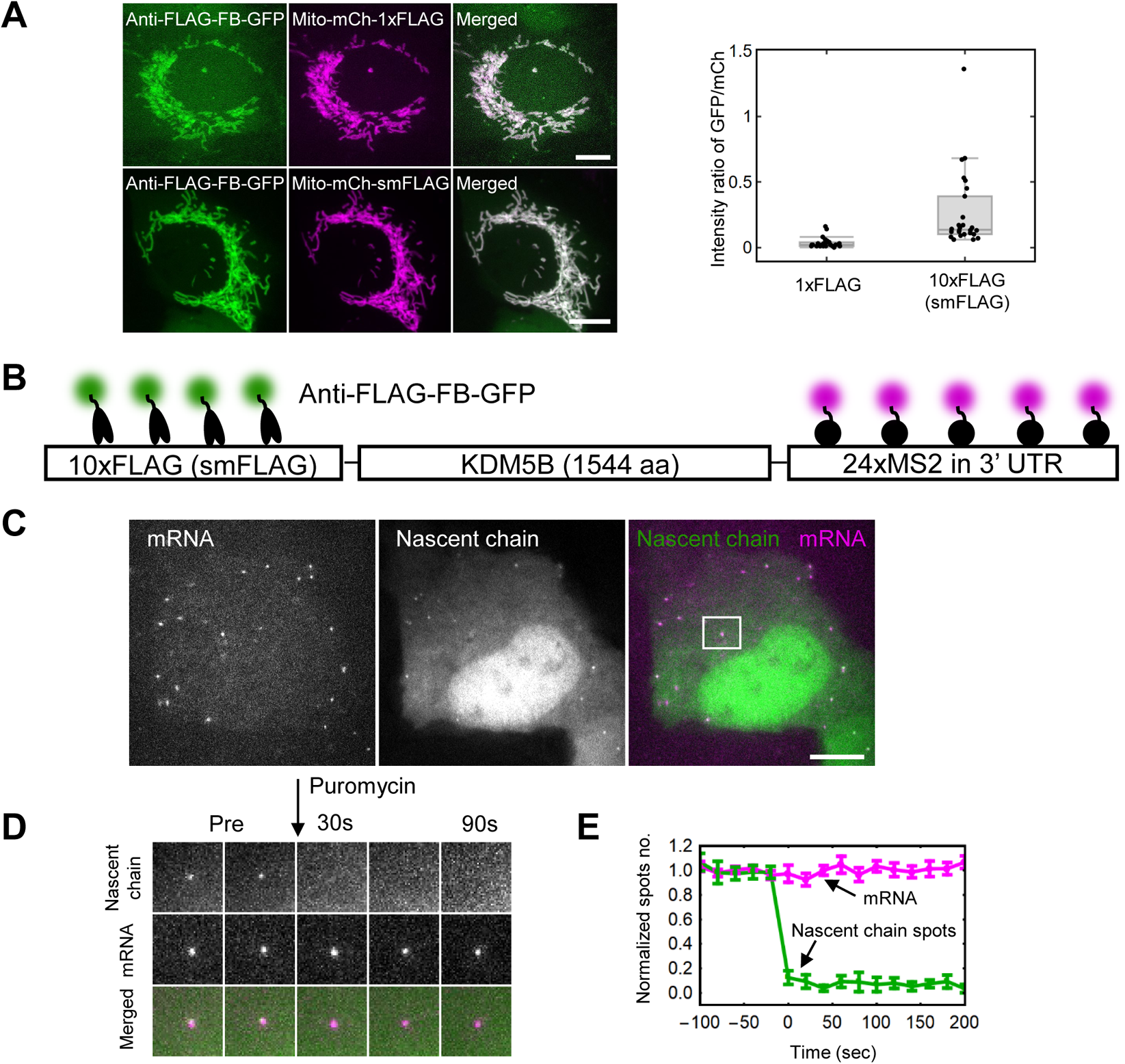
Amplifying fluorescence with the anti-FLAG frankenbody. (**A**) Left: Representative cells expressing the anti-FLAG frankenbody (green) labeling mitochondria with 1× or 10× FLAG (smFLAG) tags (magenta); Right: Box plot of the fluorescence intensity ratio of the anti-FLAG FB-GFP labeling Mito-mCh-1×FLAG or -10×FLAG (smFLAG). Center lines show the medians; the boxes indicate 25-75%; whiskers extend 1.5 times the interquartile range from the 25th and 75th percentiles. The ratio of net intensities after background subtraction in each channel is 0.032±0.007 (Mean ± SEM) for 1× FLAG (n=30 cells) and 0.250±0.050 (Mean ± SEM) for 10× FLAG (n=30 cells), indicating, on average, 7.7±2.2 (Mean ± SEM) anti-FLAG FB-GFP bind to 10× FLAG (smFLAG). (**B**) A diagram depicting frankenbody (anti-FLAG FB-GFP; green) and MCP-HaloTag/JF646 (magenta) labeling FLAG epitopes and mRNA MS2 stem loops, respectively, in a KDM5B translation reporter. (**C**) A representative cell showing the colocalization of anti-FLAG FB-GFP (green) with KDM5B mRNA (magenta). (**D**) A montage of the representative translation spot highlighted in **B** showing the disappearance of nascent chain signals within seconds of adding the translational inhibitor puromycin. (**E**) The normalized number of mRNA spots (magenta) and nascent chain spots (green) as a function of time after adding the translational inhibitor puromycin (n=9 cells in 3 independent experiments, error bars show the cell-to-cell SEM). Scale bars, 10 µm.

For tracking nascent chains with anti-FLAG frankenbody, a custom-built widefield fluorescence microscope based on a highly inclined and laminated optical sheet (HILO) illumination scheme (Tokunaga et al., 2008) was used. Briefly, the excitation beams, 488, 561, 637 nm solid-state lasers (Vortran), were coupled and focused off-axis on the rear focal plane of the objective lens (APON 60XTIRF, NA 1.49; Olympus). The emission signals were split by an imaging grade, ultra-flat dichroic mirror (T660lpxr, Chroma). The longer emission signals (far-red) after splitting were passed through a bandpass filter (FF01-731/137-25, Semrock). The shorter emission signals (red and green) after splitting were passed through either a bandpass filter for red (FF01-593/46-25, Semrock) or a bandpass filter for green (FF01-510/42-25, Semrock) installed in a filter wheel (HS-625 HSFW TTL, Finger Lakes Instrumentation). The longer (far-red) and the shorter (red and green) emission signals were detected by two separate EM-CCD cameras (iXon Ultra 888, Andor) by focusing with 300 mm achromatic doublet lenses (AC254-300-A-ML, Thorlabs). The combination of the 60× objective lens from Olympus, a 300 mm tube lens, and iXon Ultra 888 produces 100× images with 130 nm pixel^−1^. A stage top incubator for temperature (37 °C), humidity, and 5% CO_2_ (Okolab) is equipped on a piezoelectric stage (PZU-2150, Applied Scientific Instrumentation) for live cell imaging. The lasers, the cameras, the piezoelectric stage, and the filter wheel were synchronized by an open source microcontroller, Arduino Mega board (Arduino). Imaging acquisition was performed using open source Micro-Manager software (1.4.22).

The imaging size was set to the center 512 × 512 pixels^2^ (66.6 × 66.6 µm^2^), and the camera integration time was set to 53.64 ms. The readout time of the cameras from the combination of our imaging size, readout mode (30 MHz), and vertical shift speed (1.13 µs) was 23.36 ms, resulting in an imaging rate of 13 Hz (70 ms per image). Red and green signals were imaged alternatively. The emission filter position was changed during the camera readout time. To minimize the bleed-through, the far-red signal was simultaneously imaged with the green signal. To capture the whole thickness of the cytoplasm in cells, 13 z-stacks with a step size of 0.5 µm (6 µm in total) were imaged. For Fig. 2C, cells were imaged using the above described microscope setup with a 4 s interval between frames (lasers: 488 nm, 13 µW; 637 nm, 150 µW; Laser powers were measured at the back-focal plane of the objective). The displayed image was max-projected through the whole cell volume in ImageJ.

For puromycin treatments, U2OS cells seeded on 35mm MatTek chambers with 70% confluency were loaded with 1 µg of smFLAG-KDM5B-24×MS2 (Addgene #81084), 0.5 µg of anti-FLAG FB-GFP and 130 ng of purified MCP-HaloTag protein by bead loading (Cialek et al., 2021). 3 h post bead loading, imaging chambers were stained and washed as described above. The cells showing translation spots and mRNA molecules were imaged with 20 s intervals between each time point. At each time point, 13 z-slices were imaged with 0.5 µm step size. After acquiring 5 time points as pre-treated images, puromycin was added to imaging chambers at a final concentration of 0.1 mg/mL. After adding puromycin, the cells were continuously imaged at the same conditions until the translation spots disappeared.

### ZF probes design, construction and purification

The ZF probes were designed to target a pair of chromatin contact sites on chromosome 8 (chr8:126, 150, 000-126, 750, 000). Each ZF probe contains 6 zinc finger DNA binding domains engineered using *Zinc Finger Tools* (Mandell & Barbas, 2006). In this study, the ZF probes were fused with 10× HA or FLAG epitopes at the C-terminus with a linker sequence (GSAGSAAGSGEF; (Waldo et al., 1999)). The DNA encoding ZF probes were cloned into the pTrcHis vector (Thermo Fisher Scientific) and verified by nucleotide sequencing. Then the sequence-confirmed pTrcHis-ZF vectors were transformed into BL21(DE3) (Sigma). Single colonies were picked up and cultured overnight in LB medium containing 100 μg/mL ampicillin. The culture was diluted 1:500 in 500 mL LB containing 100 μg/mL ampicillin and incubated at 37°C until OD600 reached ~0.6. After the addition of IPTG (Isopropyl β-D-1 thiogalactopyranoside; Sigma; 2 mM) and ZnCl_2_ (100 μM), cells were regrown overnight at 16°C. After harvesting cells by centrifugation at 4000x*g* for 30 min at 4°C, the pellets were washed and resuspended in 5 mL lysis buffer (50 mM NaH_2_PO_4_, 200 mM NaCl, 10 mM imidazole, pH 8.0) with protease inhibitor cocktail (Nacalai Tesque). The cells were lysed by sonicating with 50% power setting for 15 min (15 s on/off) in ice water. After the centrifugation at 15,000x*g* for 30 min at 4°C, the supernatant was collected, filtered through a 0.45 μm filter, and applied to a 1 mL HisTrap HP column (GE Healthcare) using an AKAT Prime Plus (GE Healthcare) according to the programmed His-tag elution method: equilibration with 5X column volume lysis buffer, washing with 10× column volume wash buffer (50 mM NaH_2_PO_4_, 200 mM NaCl, 100 mM imidazole, pH 8.0), eluting with 4 mL 0 to 100% gradient with elution buffer (50 mM NaH_2_PO_4_, 200 mM NaCl, 500 mM imidazole, pH 8.0), and 20 mL elution buffer.

### Electrophoretic mobility shift assays

Cy3-labeled 36 nt long single-stranded oligonucleotides which contains ZF probe target sites were mixed with the unlabeled complementary oligonucleotides in TE buffer (10 mM Tris-HCl, pH 7.5, 1 mM EDTA), heated to 95°C for 5 min and cooled down at room temperature. The binding reactions were carried out in a 20 μL system (10 mM Tris-HCl, pH 7.5, 50 mM KCl, 5 mM MgCl_2_, 100 μM ZnCl2, 1 mM dithiothreitol (DTT), 0.05% Triton X-100, and 2.5% glycerol) with various concentration of ZF probes and fixed 20 nM labeled double-stranded oligonucleotides. The binding reaction were incubated at 37°C for 30 min and separated on 8% polyacrylamide gels (FUJIFILM Wako Pure Chemical) in 1× TBE buffer (89 mM Tris, 89 mM boric acid, 2 mM EDTA) at room temperature for 1 h. The fluorescence images were captured using a gel documentation system (LuminoGraph II, ATTO). The K_D_ for ZF probe was calculated using the adapted Michaelis-Menten equation:

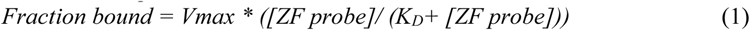

*Fraction bound* is the ratio of bound signal to the total signal (bound + unbound signal) measured by ImageJ. *Vmax* and *K_D_* were obtained by performing non-linear regression using the adapted Michaelis-Menten equation by JASP ver 0.14.1 (JASP Team; https://jasp-stats.org/).

### Live cell imaging of ZF probes

Cells were plated on a 35 mm glass-bottom dish (AGT Technology Solutions) at least a day before imaging. The medium was replaced to FluoroBrite DMEM (Thermo Fisher Scientific) containing 10% FCS and 1% GPS solution (Sigma-Aldrich) after ZF probe administration at 5 μM. This concentration was optimized to observe the highest foci-to-background ratio with substantial brightness. Fluorescent images were acquired using a spinning disk confocal microscope (Ti-E, Nikon; CSU-W1, Yokogawa Electric) equipped with a stage incubator at 37°C with 5% CO_2_ (Tokai Hit) with an EM-CCD (iXon2; Andor) and a laser system (488, 561, and 640 nm lasers; LNU-4; Nikon) using NIS-Elements ver 4.30.00 operation software (Nikon). The RAD21-GFP capture was set up as 400 ms exposure time under 75% laser transmission with 11 z-stacks (0.2 μm each). The mCherry and iRFP capture setup was adjusted based on the cell condition, typically setup as 400 to 600 ms exposure time under 100% laser transmission with 11 to 15 z-stacks (0.2 μm each). To investigate the effect of different fluorescent anti-HA frankenbody fusion tags on the appearance of foci imaged with ZF probes, wild-type HCT-116 cells were plated on 35 mm glass-bottom dishes (Mat-Tek) and transfected with expression vectors of frankenbodies tagged with monomeric EGFP (mEGFP), EGFP, mCherry, and iRFP, using the Neon Transfection system (Thermo Fisher Scientific; 0.5 μg DNA and 1× 10^5^ cells using 10 μL tip; 1,100 V; 30 ms pulse). 48 h later, ZF-F probe was administrated for 6 h before fluorescence images were acquired.

Image analysis and processing were performed using ImageJ2 (Rueden et al., 2017). Linear contrast enhancements were applied to better represent foci in individual images. The same settings were used for time-lapse images. To measure the distance between two foci labeled with ZF probes, a pair of foci in the same section or within 2 z-sections were selected for 2D analysis. Their focus areas were first defined by thresholding and then their centers of mass were determined by the ImageJ2 Measurement function. In the event that both alleles were not in close association, the distances for all combinations of foci were calculated and the nearest ones were defined as a pair and the remaining ones were defined as another pair. In time-lapse analysis, a pair could be tracked or back tracked. For Fig.6A, the area of the nucleus was selected manually by visual inspection to measure RAD21-mAID-mClover-GFP signals. Statistical analyses were performed using JASP ver 0.14.1 (JASP Team; https://jasp-stats.org/). Plots were generated using the plotly package (Plotly Technologies Inc.; https://plot.ly) in the R language (R Core Team; https://www.R-project.org/). Outliers were defined as points above or below 1.5 times the interquartile range from the 25th and 75th percentile. Fig. 3 was created using BioRender.com with a student plan subscription.

**Fig. 3.**
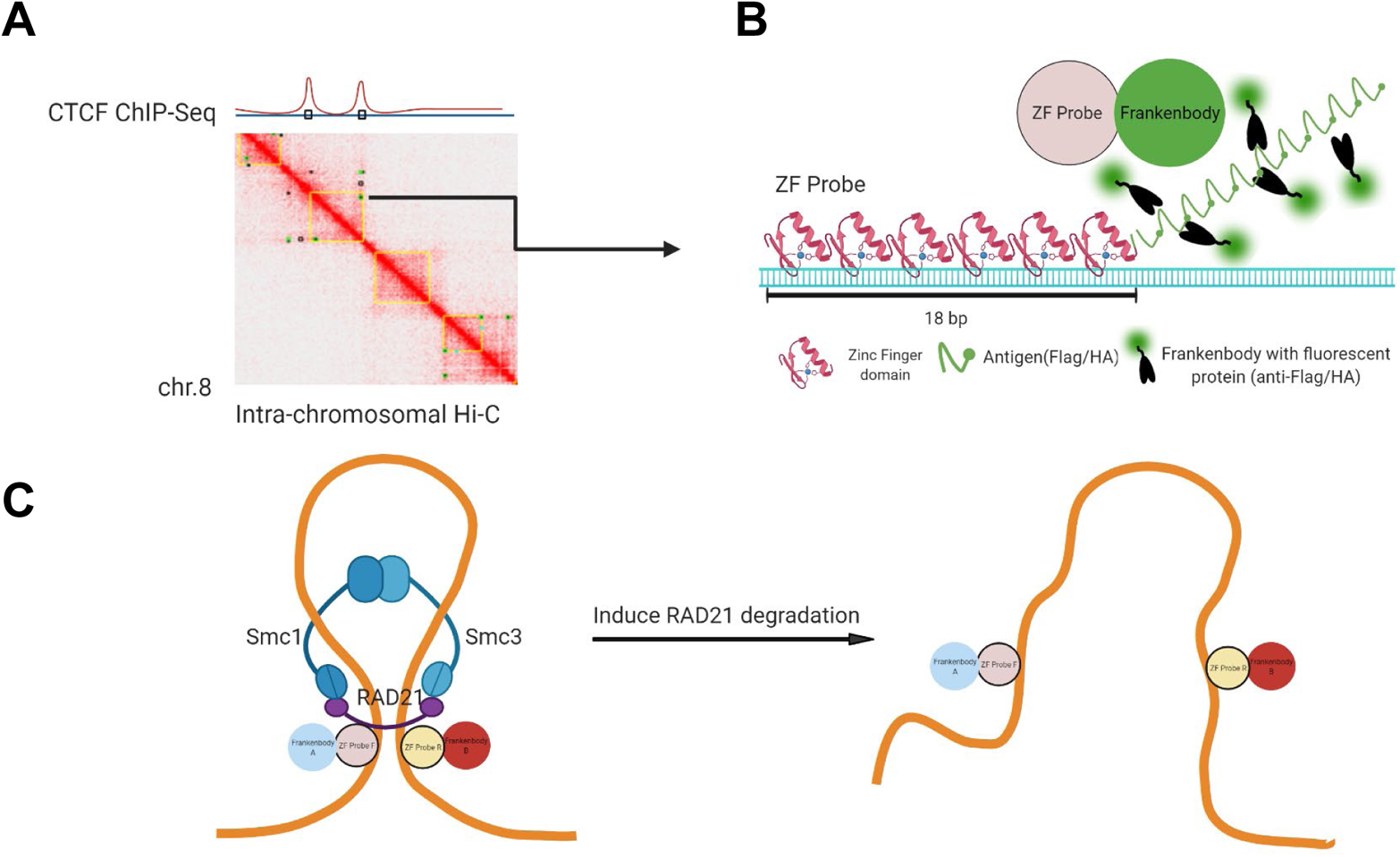
Designing a pair of zinc-finger protein probes to track chromatin contact sites. (**A**) Chromatin contact sites predicted by different HiC algorithms are chosen. (**B**) A schematic of a DNA-bound ZF probe that contains 6 DNA recognition domains and 10× HA or FLAG linear epitopes for signal amplification upon binding of complementary fluorescent frankenbodies. (**C**) A schematic of chromatin looping and unlooping. Two ZF probes that bind to the sites of chromatin contact are expected to be observed within a close distance of one another (D_C_; the probe distance in the presence of a loop). When cohesin is absent, the two probes will be more separated (D_L_; the probe distance in the absence of a loop).

**Fig. 6.**
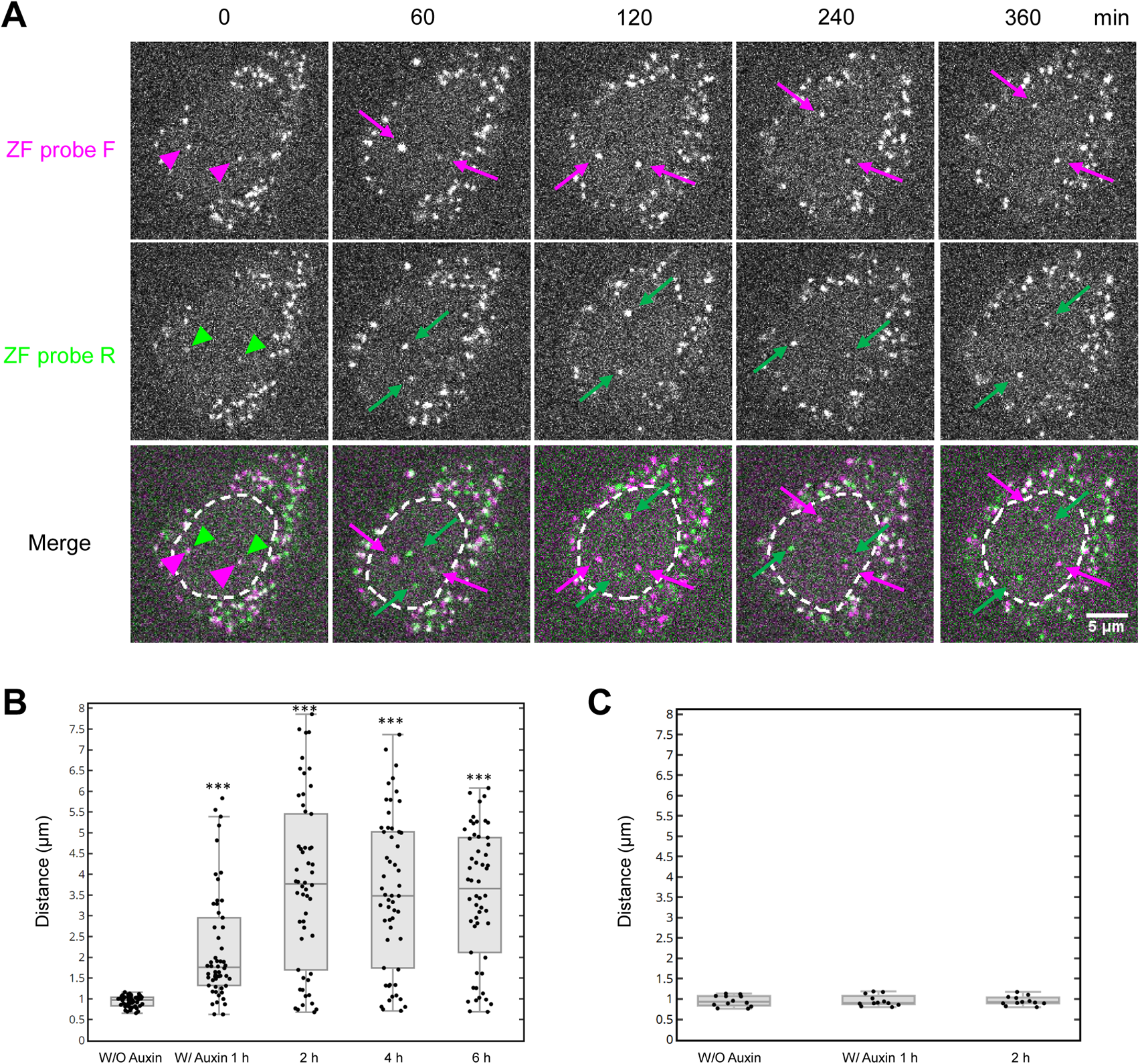
Dissociation of genomic loci induced by cohesin degradation. HCT116-mAID-RAD21-mClover cells were transfected with anti-FLAG and anti-HA frankenbody expression vectors, loaded with ZF probes, and were treated (**A** and **B**) or untreated (**C**) with auxin. (**A**) Representative confocal images of living HCT116-RAD21-mAID-mClover cells loaded with ZF probe F (mCherry frankenbody; red) and R (iRFP frankenbody; green). The nuclear foci of ZF probes F and R were closely associated 5 min after addition of auxin (red and green arrowheads), but became separated in 30 min (magenta and green arrows). Nuclear areas are indicated in dashed line in merged images. Scale bar, 5 μm. (**B**) Box plot of the distance between ZF probes F and R at the indicated time point after auxin addition. ZF probe pairs that were originally associated (< 1.2 μm) before auxin addition (W/O Auxin) were selected and tracked (n=54 pairs from 2 dishes). The statistical analysis was performed between W/O Auxin and others by Wilcoxon signed-rank test (*** p <0.001; see Supplementary Table S5). (**C**) Box plot of the distance between ZF probes F and R in mock treated cells without auxin (n=12 from 2 dishes). There is no significant difference between the different time points by Wilcoxon signed-rank test. For box plots, center lines show the medians; the boxes indicate 25-75%; whiskers extend 1.5 times the interquartile range from the 25th and 75th percentiles.

### Estimating the number of fluorescence molecules in the foci

The fluorescence intensity of the foci and single EGFP molecules were measured on a custom-build inverted microscope IX83 (Olympus) equipped with a 100× objective lens (UPlanSApo NA 1.40; Olympus) and a 2× intermediate magnification lens (Olympus) (Lim et al., 2018). HILO illumination (Tokunaga et al., 2008) of a 488-nm laser (OBIS 488 LS; Coherent; 100 mW) was used with the CellTIRF system (Olympus). Images were acquired at an exposure time of 32.55 ms using an EM-CCD (C9100-13; Hamamatsu Photonics). The EM gain was set to 180 to avoid the saturation of the foci intensity, and the two frames were averaged to increase the signal-noise ratio of the single-molecule intensity. The fluorescence intensity was measured using ImageJ software. The single-molecule intensity was determined using a circular area with a 7-pixel diameter corresponding to the size of the Airy disk, and the outer circular region with 2-pixel thickness was used as the background. To convert the intensity of the foci into the number of molecules, the total intensities of the foci from 45 cells were divided by the averaged single-molecule fluorescence from 5 extremely low-expressing cells.

### Lentivirus-mediated knockdown by shRNA expression

The addgene pLKO.1 lentiviral knockdown protocol was adapted. Gene-specific shRNA and scrambled shRNA were designed using the TRC shRNA Design online tool on GPP Web Portal (https://portals.broadinstitute.org/gpp/public/; Yang et al., 2011; Supplementary Table S1). HEK-293T cells were grown up to ~80% confluency in a 6 cm culture dish and were transfected with psPAX2, pCMV-VSV-G, and pLKO.1 puro-based plasmid harboring the designed shRNA (pCMV-VSV-G and pLKO.1 puro were a gift from Bob Weinberg, Addgene plasmid #8454; http://n2t.net/addgene:8454; RRID: Addgene_8454; psPAX2 was a gift from Didier Trono, Addgene plasmid #12260; http://n2t.net/addgene:12260; RRID: Addgene_12260) using Lipofectamine 3000 (Thermo Fisher Scientific) following the manufacturer’s instructions. One day after the transfection, the medium was replaced with 5 ml fresh DMEM, which was harvested the next day. The harvested medium (6 mL) was filtered through a 0.2 μm filter and added to HCT116 cells at ~70% confluent in a 6 cm dish (5 mL) with 8 μg/mL polybrene. After an incubation period of 24 h, the medium was replaced to fresh DMEM with 2 μg/mL puromycin for at least 3 days to select infected cells before analysis.

The efficiency of shRNA-mediated knockdown was assessed by immunoblotting. Cells were washed with ice-cold PBS twice and lysed by RIPA buffer (50 mM Tris-HCl, pH 8.0, with 150 mM sodium chloride, 1.0% Igepal CA-630 (NP-40), 0.5% sodium deoxycholate, and 0.1% sodium dodecyl sulfate) at 1× 10^7^/ml. After denaturation by heating at 95°C for 10 min, the protein concentration was measured using a bicinchoninic acid assay kit (FUJIFILM Wako Pure

Chemical) and adjusted to 2 mg/mL. Cell lysates (10 μg each) were separated on 7.5% polyacrylamide gels and transferred on to PVDF membranes (Pall). Then the membranes were incubated in Blocking One (Nacalai Tesque) for 30 min, incubated with the primary antibody in Can Get Signal Solution I (TOYOBO) overnight at 4°C with gentle shaking, washed in TBS-T (20 mM Tris-HCl, pH 8.0, 150 mM NaCl, 0.05% Tween 20) for 10 min three times, and incubated with the secondary antibody for 1 h at room temperature in Can Get Signal Solution II (TOYOBO). After washing with TBS-T for 10 min three times, the membranes were incubated with ImmunoStar LD (FUJIFILM Wako Pure Chemical) for 5 min before detecting the chemiluminescent signals using a LuminoGraph II gel documentation system (ATTO).

## RESULTS AND DISCUSSION

### Development and evaluation of an anti-FLAG frankenbody to image FLAG epitopes in living cells

To label and track a pair of endogenous genomic loci in single living cells with minimal perturbation, we envisioned developing a pair of complementary ZF-based probes, the first harboring a 10× HA tag and the second a 10× FLAG tag for signal amplification in two separate colors. To amplify signals at the 10× HA tag, we recently developed a chimeric anti-HA single-chain variable fragment (scFv), which we refer to as the anti-HA “frankenbody” (Zhao et al., 2019). To similarly amplify signals at the 10× FLAG tag, we set out to develop a complementary anti-FLAG frankenbody (Fig. 1A; Supplementary Fig. 1A).

We began with an anti-FLAG scFv generated from a monoclonal antibody against the DYKDDDDK tag. Unfortunately, this wild-type anti-FLAG scFv (wtFLAG-scFv) did not show any binding affinity to the FLAG tag in living U2OS cells. In particular, a GFP-tagged version of the wtFLAG-scFv did not colocalize in the cell nucleus with a FLAG-tagged mCherry-H2B (Supplementary Fig. 1B and 1C). We hypothesized that the loss of binding affinity is due to misfolding of the scFv. This is a common issue for scFvs since their heavy and light chains are fused in an unnatural way and their disulfide bonds are reduced in the cytoplasm of living cells (Ewert et al., 2004; Zhao et al., 2019).

To overcome these issues, we engineered more soluble chimeric scFv frankenbodies by grafting all six complementary determining regions (CDRs) or loops of the wtFLAG-scFv to scFv scaffolds that have already been demonstrated to fold correctly in living cells (Fig. 1A; Supplementary Fig. 1A). We chose five scFv scaffolds as loop-grafting candidates: 1) an scFv that binds to histone H4 mono-methylated at Lysine 20 (H4K20me; 15F11; Sato et al., 2016); 2) an H4K20me2-specific scFv (2E2; Zhao et al., 2019); 3) an H3K9ac-specific scFv (13C7; Sato et al., 2013); 4) a Suntag-specific scFv (Tanenbaum et al., 2014); and 5) a bone Gla protein (BGP)-specific scFv (KTM219) (Wongso et al., 2017). Among these five scFv scaffolds, their sequence identity with wtFLAG-scFv ranged from 43-74% for the variable region of the heavy chain (VH) and from 58-82% for the variable region of light chain (VL), as shown in Supplementary Table S2.

In general, the higher the sequence identity in the VH and VL regions, the more likely loop grafting will be successful (Ewert et al., 2004; Zhao et al., 2019). Among the five selected scFv scaffold candidates, 15F11 and 2E2 share the highest sequence identity with wtFLAG-scFv. We therefore predicted that grafting onto the 15F11 and 2E2 scaffolds would have a higher chance of producing a functional anti-FLAG frankenbody. As shown in Supplementary Fig. 1D and 1F, this was indeed the case. In particular, GFP-tagged frankenbodies generated by loop grafting onto the 15F11 and 2E2 scaffolds showed excellent colocalization in the nucleus with FLAG-tagged mCherry-H2B, while the other frankenbodies did not colocalize in the nucleus. Moreover, in the absence of FLAG tags, the 15F11 and 2E2 anti-FLAG frankenbodies were evenly spread throughout cells (Supplementary Fig. 1E and 1F), demonstrating these frankenbodies do not have any off-target binding sites. Together, these two anti-FLAG frankenbodies complement our previously engineered anti-HA frankenbodies (Zhao et al., 2019), demonstrating the power of one-step loop grafting for generating functional intrabodies against linear epitope tags in a fast manner.

To further characterize the 15F11 anti-FLAG frankenbody, we imaged it in diverse live-cell environments. For this, we co-transfected the monomeric enhanced GFP-tagged frankenbody (anti-FLAG FB-mEGFP) with plasmids encoding various FLAG-tagged proteins, including the nuclear protein H2B (4×FLAG-mCh-H2B, also used in the initial screen in Supplementary Fig. 1D), the cytoplasmic protein β-actin (4×FLAG-mCh-β-actin), and the mitochondrial protein mitoNEET (Mito-mCh-1×FLAG; Colca et al., 2004). In all cases, the anti-FLAG FB-mEGFP was highly co-localized with the FLAG-tagged proteins (Fig. 1B). In particular, plots of the fluorescence in the two channels along lines spanning each cell revealed high correlations with Pearson correlation coefficients all greater than 0.98 (Fig. 1C). These data demonstrate that the anti-FLAG frankenbody can bind specifically to the FLAG tag fused to both the N- and C-terminal ends of proteins in diverse cellular settings.

We next characterized the functionality of the anti-FLAG frankenbody when fused to different tags beyond mEGFP, including mRuby2, mCherry, SNAP-tag, and HaloTag. To do this, we co-transfected into U2OS cells a plasmid encoding the anti-FLAG frankenbody fused to each fluorescent protein along with a plasmid encoding FLAG-tagged H2B. All exhibited H2B-like distributions in nuclei. After imaging, we approximated the labeling efficiency of the frankenbody fusions by calculating the ratio of the intensity of frankenbody fluorescence in the cell nucleus (where FLAG-tagged H2B localizes) to the cytoplasm (where there is presumably very few FLAG tags) (Fig. 1D). This revealed the labeling efficiency of the frankenbody is similar when fused with SNAP-tag, HaloTag and mEGFP (p-value > 0.05). Interestingly, fusions to mCherry tended to aggregate (data not shown), whereas fusions to the other red fluorescent protein, mRuby2, actually had the highest labeling efficiency (p-value < 0.05). Thus, these data demonstrate the anti-FLAG frankenbody can work when fused to a variety of complementary fluorescent proteins for imaging in up to four colors (e.g., by using SNAP-Cell 430 and HaloTag JF646 ligands, in addition to green and red fluorescent proteins).

Finally, we wanted to test the binding turnover time of the anti-FLAG frankenbody. For this, we performed fluorescence recovery after photobleaching (FRAP) experiments in living U2OS cells that co-expressed the anti-FLAG FB-mEGFP along with 4×FLAG-mCh-H2B. In these experiments, since H2B has a FRAP recovery time on the order of hours, any fluorescence recovery observed in the frankenbody channel that occurs on the minutes or seconds timescales can be attributed almost entirely to frankenbody turnover rather than H2B turnover. As shown in Fig, 1E, the FRAP recovery halftime (*t_half_*) of the anti-FLAG frankenbody is 35.3 ± 2.2 s. While this recovery time is faster than that of the anti-HA frankenbody (*t_half_* = 141±7 s) (Zhao et al., 2019), the timescale is longer than other scFv we have developed to image histone modifications (Sato et al., 2013). Thus, although binding might not be as tight as the anti-HA frankenbody, the more rapid turnover can be advantageous in certain instances; for example, rapid turnover can minimize interference with tagged protein functionality and also minimize photobleaching (since frankenbody photobleached in the imaging plane can unbind and be replaced by a non-photobleached frankenbody).

### Signal amplification using the anti-FLAG frankenbody

Before coupling the anti-FLAG frankenbody with ZF proteins for genomic loci visualization, we first wanted to verify it could be used for signal amplification. To test this, we coexpressed anti-FLAG FB-mEGFP along with mito-mCh fused either to a 10× FLAG spaghetti monster tag (mito-mCh-smFLAG) or a 1× FLAG tag (mito-mCh-1×FLAG). To gauge signal amplification, we imaged the two combinations of plasmids under the same conditions and quantified the ratio of background-subtracted GFP to mCherry fluorescence at mitochondria (Fig. 2A). For the mito-mCh-1×FLAG construct, the ratio was 0.032 ± 0.007 (Mean ± SEM; n=30), while for the mito-mCh-smFLAG construct, the ratio was significantly higher at 0.250 ± 0.050 (Mean ± SEM; n=30). Dividing the ratios implies there are on average 7.7 ± 2.2 (Mean ± SEM) more anti-FLAG frankenbodies binding to smFLAG than to 1× FLAG, suggesting significant signal amplification.

To further test the limits of signal amplification, we next used the anti-FLAG frankenbody to track the translation dynamics of single reporter mRNAs using Nascent Chain Tracking (Morisaki et al., 2016). For this, we transfected cells with a reporter plasmid encoding 24×MS2 stem loops in the 3’UTR and a 10× smFLAG at the N-terminus of the nuclear protein KDM5B (Fig. 2B). With this reporter, single-mRNA translation sites can be detected by the colocalization of reporter mRNA (labeled by bead-loaded Halo/JF646-tagged MS2 coat proteins) and nascent protein (labeled by co-expressed anti-FLAG FB-mEGFP). As shown in Fig. 2C and 2D, we could indeed detect and track multiple single mRNA translation sites, and each site was sensitive to the addition of the translation inhibitor puromycin. Thus, the anti-FLAG frankenbody can amplify signals from 10× FLAG tags for single molecule tracking experiments.

### Design, expression, purification, and evaluation of Zinc-finger DNA-binding protein probes

With the anti-FLAG and anti-HA frankenbodies in hand, we next wanted to couple them with complementary ZF probes to mark and track a pair of genomic loci in living cells in two separate colors. With this goal in mind, we engineered a pair of ZF probes to precisely bind two closely-associated genomic regions in 3D cellular space. In the design process, we followed three guiding principles. First, the ZF target sequence should be within 10 kb from the chromatin contact sites considering the resolution of light microscopy (Banigan et al., 2020; Brandão et al., 2021). Second, because ZF probes have a strong binding affinity to target DNA, albeit weaker than dCas9 with sgRNA, the ZF target sequence should not overlap the functional binding motifs of common DNA-binding regulators, CpG islands, or any sequence that has the possibility for high DNA methylation. Third, non-existing ZF binding domain sequence combinations should be avoided, as each ZF domain recognizes 3 bp of DNA and the available target triplet sequences are limited (Durai et al., 2005).

To choose a model region of chromatin contact sites, we analyzed the GSE104334 data set, which is a HiC contact map of the HCT116-AID system cell line (Rao et al., 2017). We used HiC-Pro (Servant et al., 2015) and Juicebox (Robinson et al., 2018) and aligned with HCT116 Hi-C (Li et al., 2012), ENCODE HCT116 CTCF ChIP-Seq (ENCSR000BSE), and RNA-Seq (ENCSR000CWM) data (Supplementary Fig. S2). We picked a pair of genomic loci predicted by both algorithms to be in frequent contact on chromosome 8 (Fig. 3A and Supplementary Fig. S2). We used Zinc Finger Tools (Mandell & Barbas, 2006) to generate DNA sequences encoding two ZF probes that bind to the target sequences 600 kb apart. This resulted in a pair of ZF probes, each containing 6 ZF binding domains to recognize 18 bp of unmethylated DNA. Finally, we tagged each ZF probe F (ZF-F) with a 10× linear HA tag and each ZF probe R (ZF-R) with a 10× linear FLAG tag to efficiently recruit anti-HA and FLAG frankenbodies for two-color signal amplification (Fig. 3B). We reasoned this system would allow us to test if the target genomic loci become separated upon auxin-induced RAD21 degradation in living HCT116-RAD21-mAID-mClover cells (Fig. 3C).

We next expressed and purified His-tagged ZF probes in *E. coli* (Supplementary Fig. S3). We evaluated the binding specificity and affinity of the purified ZF probes (ZF-F and ZF-R) using an electrophoretic mobility shift assay and Cy3-labeled double-stranded oligonucleotides. The mobility of oligo-DNA containing the ZF target sequence was shifted in the presence of the corresponding ZF probes, whereas those with a few nucleotide substitutions did not show mobility shifts (Fig. 4A and 4B). These data indicate that the ZF probes selectively bind to the target sequences, as designed. We also estimated the binding affinity of the ZF probes to oligo-DNA, by varying protein concentrations (1-150 nM) with fixed concentrations of oligo-DNA (15 nM) in the assay. The dissociation constant (K_D_), calculated by the adapted Michaelis-Menten equation, was 42 ± 3 and 41 ± 3 nM (Mean ± SEM) for ZF probes F and R, respectively (Fig. 4C and 4D). These dissociation constants are higher than eukaryotic DNA binding proteins (K_D_ μM level), but lower than typical bacterial repressors (K_D_ pM level) (23, 24) and sgRNA-dCas9 (K_D_ a few nM) (Cencic et al., 2014; Jiang et al., 2015).

**Fig. 4.**
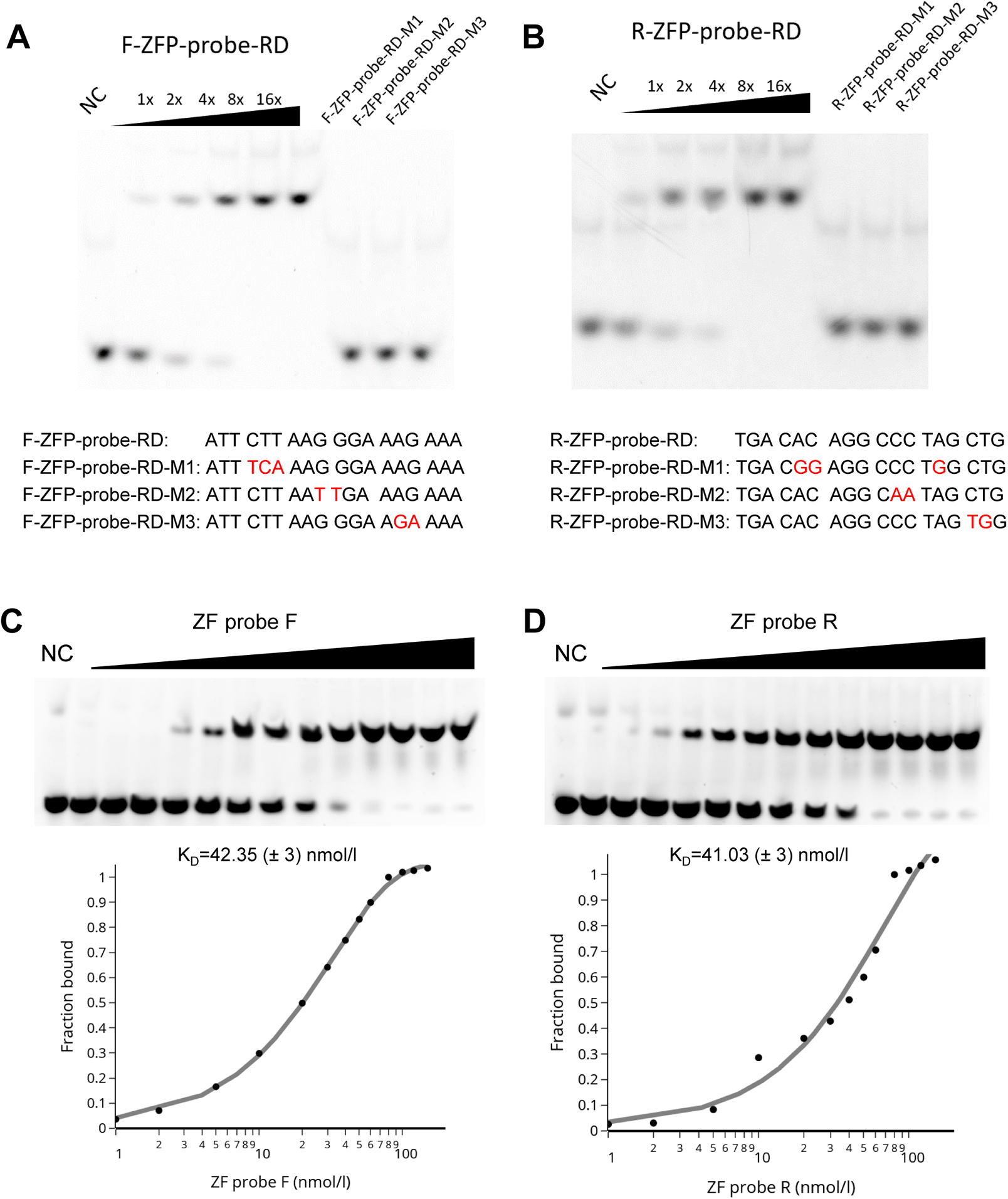
Binding specificity and affinity of ZF probes. Binding of ZF probes with DNA was analyzed by a gel mobility shift assay. (**A** and **B**) Double stranded Cy3-labeled DNA (20 nM) was mixed with ZF probes F (**A**) and R (**B**) (6.25, 12.5, 25, 50, and 100 nM for DNA containing the target sequence; and 100 nM for control DNA). Oligonucleotide sequences and the mutation sites (in red) are indicated. ZF probes bound specifically to the target sequence. (**C** and **D**) Double stranded Cy3-labeled DNA containing the target sequence (20 nM) was mixed with ZF probes F (**C**) and R (**D**) (1-150 nM). The binding affinity (dissociation constant; K_D_) was calculated based on the curve of bound fractions of DNA with respect to the ZF probe concentration.

### ZF probes were closely associated depending on cohesin in living cells

Having tested the ZF probes *in vitro*, we next tested their ability to label endogenous DNA in living HCT116-RAD21-mAID-mClover cells (Natsume et al., 2016). In these cells, a cohesin subunit RAD21 gene was replaced with RAD21-mAID-mClover, the degradation of which can be induced by auxin (Fig. 5A). HCT116-RAD21-mAID-mClover cells were transfected with expression plasmids for anti-HA and anti-FLAG frankenbodies (anti-HA mCherry-FB and anti-FLAG iRFP-FB). 48-96 h after transfection, purified ZF probes were added in the medium at a final concentration of 5 μM each for 2 h, during which time the ZF probes were incorporated into cells. Without ZF probes, transiently expressed frankenbodies distributed throughout the nucleus and cytoplasm (Fig. 5B). Although cytoplasmic aggregations were often observed in this cell line particularly under a high laser excitation that is needed to detect weak ZF probe signals, no nuclear foci were formed without ZF probes (Fig. 5B, top). By contrast, in cells incubated with ZF probes, two nuclear foci were observed (Fig. 5B, bottom). Some ZF-F and ZF-R foci were close together (arrowheads) and others were far apart (arrows).

**Fig. 5.**
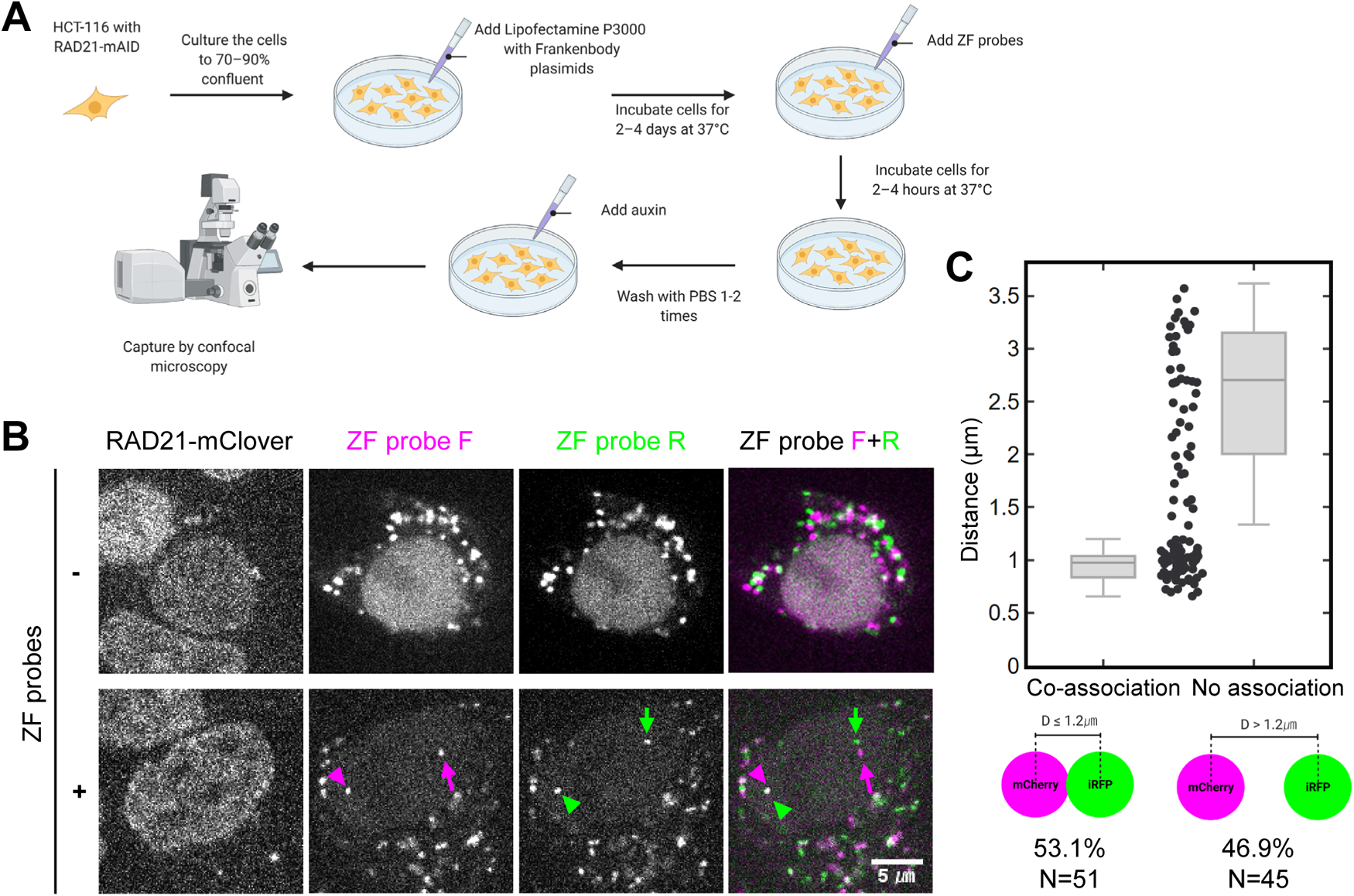
Visualizing genomic foci using ZF probes and frankenbodies in living cells. (**A**) A flow chart of the genome visualization method. Cells are transfected with expression vectors of anti-HA mCherry-frankenbody (FB) and anti-FLAG iRFP-FB and then incubated with the HA- and Flag-tagged ZF probes, which are incorporated into cells. (**B**) Representative single confocal sections of living HCT116-RAD21-mAID-GFP cells that express frankenbodies without (-) and with (+) incubation with ZF probes. Frankenbodies are seen in the cytoplasm without ZF probes, but two nuclear foci in the nucleus (arrowheads and arrows) were only observed after incubation of cells with ZF probes F (magenta) and R (green). In the nucleus with ZF probes, one pair of probes were located close together (arrowheads) and another pair were far apart (arrows). (**C**) Distance between the nearest ZF probes F and R. The distances of all pairs (96 from 3 dishes) are plotted in the middle. As there is a gap at a separation distance of ~1.2 μm and separation distances as large as 3.5 μm were observed, a threshold was set at 1.2 μm to classify probe pairs into two groups with co-association (53%; 51 out of 96) or no association (47%; 45 out of 96). Scale bar, 5 μm.

In principle, if a single ZF probe harboring 10 epitopes is localized to a target genomic locus, up to 10 fluorescent frankenbodies can accumulate. However, the fluorescence signals of nuclear foci we observed were brighter than expected, suggesting more than one ZF probe per locus. We speculated that fluorescent protein-mediated multimerization and/or aggregation could bridge multiple ZF probes (Supplementary Fig. S4A).

To test this possibility, we compared the fluorescence enrichments in foci using anti-HA frankenbody tagged with different fluorescent fusion tags, including monomeric EGFP (mEGFP), EGFP, mCherry, and iRFP. mEGFP has a mutation at the dimerization surface of EGFP that eliminates nearly all dimerization (Zacharias et al., 2002), while EGFP is known to weakly dimerize (Zacharias et al., 2002) and iRFP forms a stable dimer (Filonov et al., 2011). mCherry is known to be a monomeric protein, but when fused with other proteins, including the anti-HA frankenbody, mCherry-tagged proteins can aggregate, probably by multimerization (Landgraf et al., 2012). In each experiment, we loaded ZF-F probes two days after transfecting the frankenbody expression vectors into wild-type HCT116 cells. Nuclear foci were clearly observed in cells expressing anti-HA frankenbody tagged with EGFP, mCherry, and iRFP, but the foci with anti-HA frankenbody tagged with mEGFP were only ambiguously detected (Supplementary Fig. S4B and S4C; Supplementary Table S3). These data support the notion that fluorescent protein-mediated multimerization can lead to the formation of bright foci that contain many ZF probes and frankenbodies.

By comparing the intensity of the foci with that of single molecule EGFP (Joglekar et al., 2008), we determined a focus contained on average ~140 EGFP molecules (Supplementary Fig. S4D; Supplementary Table S4). Therefore, around ten (or more) ZF probes and hundreds of frankenbodies appear to accumulate in a nuclear focus (Supplementary Fig. S4A). Presumably, it is the binding of a single ZF probe to target DNA that seeds the formation of such a large complex because we only observed a few bright foci per nuclei. Once a sufficiently large complex stochastically forms (which can take time), its binding to target DNA becomes significantly more stable than any single ZF probe (which binds with ~40 nM affinity), most likely because the large number of constituent ZF probes within the complex facilitate very rapid rebinding following an unbinding event.

Assuming the bright ZF-F and ZF-R foci we observed did indeed mark target DNA, we next measured the distance between them. We found neighboring foci were anywhere from 0.6 to 3.7 μm apart (n=96) (Fig. 5C). As clusters from 0.6-1.2 μm were observed, we set 1.2 μm as a co-association cut-off point to categorize the spot distance into co- and no-association groups (Fig. 5C). Although this 1.2 μm cut-off distance appears to be larger than the typical distance of contact sites observed by high resolution FISH analyses, the distance of two genomic sites can be highly heterogenous (Finn et al., 2019; Rao et al., 2014). To simplify matters, we conservatively used the cut-off to simply classify spots, rather than pinpoint their true physical separation. Among 96 pairs, 51 (53.1%) were co-associated according to our cut-off (Fig. 5C). To validate the specificity of the ZF probes for target DNA, we used a mouse cell line that contained the target sequence to the ZF-F probe, but not the ZF-R probe. In these cells, as expected, ZF-F, but not ZF-R, showed foci in mouse cells that were brightly labeled by frankenbodies (Supplementary Fig. S5). Time-lapse imaging revealed that the ZF probe accumulated in foci in the nucleus within 1 h (Supplementary Fig. S4A).

To analyze the contribution of cohesin on the association of the two ZF probes, cohesion degradation was induced by auxin in HCT116-RAD21-mAID-mClover cells (Natsume et al., 2016). After the addition of auxin, the overall RAD21-mClover signal was decreased to 15% in 2 h and further down to 7% in 4 h (Supplementary Fig. S6), confirming that the auxin-induced degradation system was effective as reported (Natsume et al., 2016). Cells expressing frankenbodies were loaded with the ZF probes and fluorescence images were captured every 1 h immediately after auxin addition (Fig. 6A). To monitor the effect of cohesin depletion, we tracked pairs of ZF-F and ZF-R foci that were originally < 1.2 μm apart (Fig. 6B; the first column). Those pairs became substantially more separated in 1 h and afterwards (Fig. 6B). Analysis of 54 pairs of ZF probes revealed that their distances were significantly far apart in 1 h and reached steady state separation in 2 h (Fig. 6B; Supplementary Table S5). These data are consistent with the defect of cohesion induced by RAD21 degradation (Rao et al., 2017) (Fig. 3C). In control cells without auxin, the distance of the ZF probes remained unchanged for 2 h (Fig. 6C).

We next analyzed the function of CTCF and WAPL (Wings apart-like protein homolog) in the association of ZF-F and ZF-R foci by knockdown, both individually and simultaneously using a lentivirus shRNA expression system (Fig. 7A). CTCF is a zinc-finger protein that can act as an anchor to define the boundary for CTCF-dependent cohesin (de Wit et al., 2015; Parelho et al., 2008; Rao et al., 2017) and WAPL regulates cohesin dynamics by disassembling the cohesin ring complex (Gerlich et al., 2006; Kueng et al., 2006; Tedeschi et al., 2013). Knockdown of both CTCF and WAPL results in loosening chromatin contacts. This is because CTCF depletion disrupts cohesin-anchoring at specific sites and WAPL depletion leads to an extended loop formation that increases the size of topologically associating domains (Wutz et al., 2017). Immunoblotting showed the levels of CTCF and WAPL were decreased to roughly 17% and 6%, respectively, by single shRNA expression, and 22% and 8%, respectively, by double expression (Fig. 7A). We measured the distance between ZF-F and ZF-R foci in knockdown cells that were still viable under these conditions (Fig. 7B). CTCF knockdown caused the separation of the two ZF foci, but to a lesser degree than RAD21 degradation (Fig. 7C). WAPL knockdown had a milder effect on the probe distance compared to CTCF knockdown (Fig. 7C). In WAPL knockdown, the number of well separated pairs decreased, indicating that 75% of the pairs were less than ~2 μm apart compared to the scramble controls in which 75% were less than ~2.6 μm apart (Fig. 7C). This global shortening of genomic foci is consistent with more loop formation due to a lack of cohesin remover (Wutz et al., 2017). The double knockdown of CTCF and WAPL appeared to have an additive effect compared to the single knockdowns, causing the separation of the two ZF foci at a similar level to cohesin degradation (Fig. 7C).

**Fig. 7.**
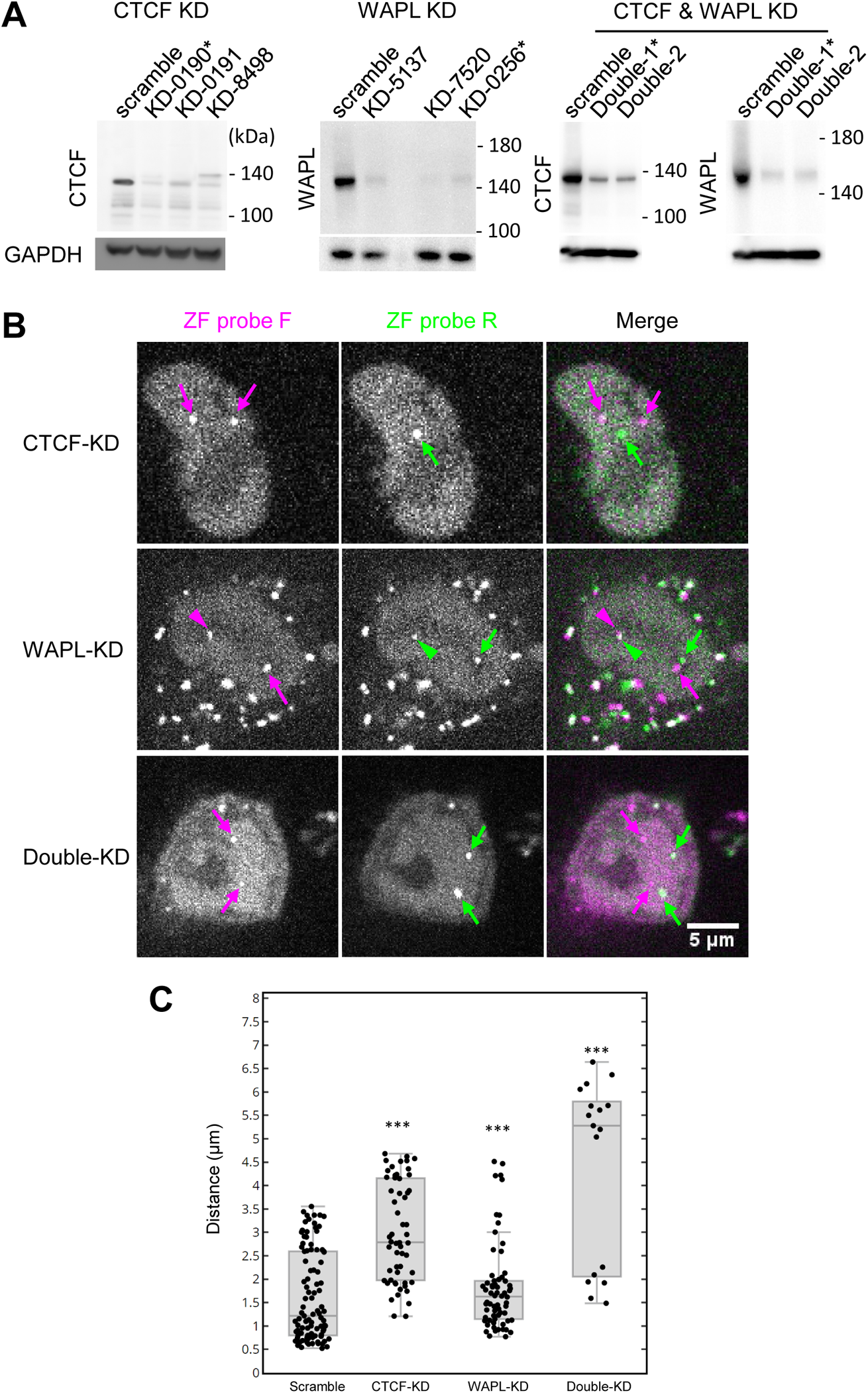
Effects of CTCF and WAPL knockdown (KD) on the association of ZF probe target loci. CTCF and WAPL in HCT116-RAD21-mAID-mClover cells were independently and simultaneously knocked down by lentivirus-mediated shRNA expression. (**A**) Immunoblotting of CTCF and WAPL in KD cells. (**B**) Confocal images of ZF probes F (mCherry; magenta) and R (iRFP; green) in KD cells. Arrows and arrowheads indicate the separate and associated foci, respectively. Scale bar, 5 μm. (**C**) Box plot of ZF probe distance in control (Scramble; n=99), CTCF-KD (n=56), WAPL-KD (n=66) and CTCF and WAPL double KD cells (n=17), each from 2 dishes. The significance to “Scramble” control was analyzed by Wilcoxon signed-rank test (*** p<0.001). For box plots, center lines show the medians; the boxes indicate 25-75%; whiskers extend 1.5 times the interquartile range from the 25th and 75th percentiles.

### Genome organization was partially recovered by cohesion restoration

We finally examined if ZF foci separated by RAD21 degradation become re-associated by cohesin restoration. HCT116-RAD21-mAID-mClover cells were incubated with auxin for 4 h to degrade RAD21, and then cells were further incubated with a fresh auxin-free medium to restore the cohesin complex. The level of RAD21-mAID-mClover recovered to ~74% (n=107 cells) compared to the control without auxin treatments in 6 h (Supplementary Fig. S7A). Cell viability dropped to ~65% in auxin for 4 h and was restored to ~75% in 6-24 h after washing away auxin (Supplementary Fig. S7B). To look for signs of ZF-F and ZF-R foci re-association, we tracked pairs of ZF foci for 12 h from immediately after auxin removal (Fig. 8A). Among 22 pairs, two were still associated at a distance <1.2 μm after auxin treatment (Fig. 8B). The other 20 pairs showed significant distance reduction after 2 h RAD21 recovery (Fig. 8B and 8C; Supplementary Table S6). Even though 15 pairs with a distance >3.6 μm did not restore their co-association status, 5 pairs with a distance between 1.2 and 3.6 μm dropped below the 1.2 μm association cut-off point after a 6 h recovery (Fig. 8B). It is possible that the pairs with a distance >3.6 μm were not associated even before auxin addition, as nearly half of ZF pairs were not associated in auxin-untreated cells (Fig. 5C). However, the pairs that became co-associated were still smaller (31.8%; 7 out of 22) compared to the control (in which 53.1% were co-associated; Fig. 5C).

**Fig. 8.**
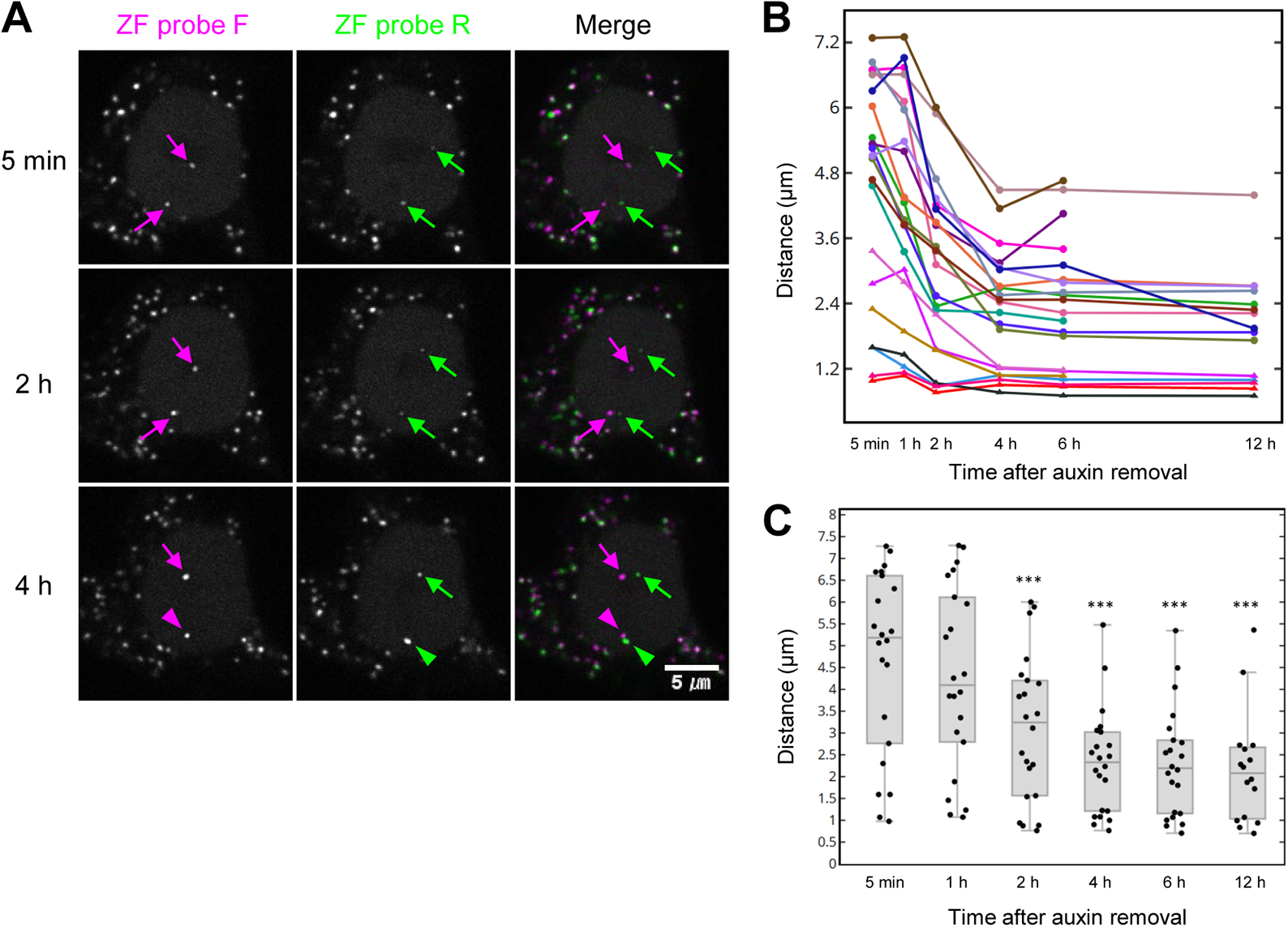
Reassociation of ZF probe target loci by RAD21 restoration. HCT116-RAD21-mAID-mClover cells were transfected with anti-FLAG and anti-HA frankenbody expression vectors, loaded with ZF probes, and treated with auxin for 4 h to degrade RAD21. After washing away auxin, cells were placed on a confocal microscope and further incubated with auxin-free medium to measure the distance between the two ZF probes. (**A**) Confocal images of ZF probes F (mCherry; Red) and R (iRPF; Green) in RAD21 recovering cells. Time after auxin removal are indicated. Arrows and arrowheads indicate the separate and associated foci, respectively. Scale bar, 5 μm. (**B**) Distances of 22 pairs of ZF probes (from 2 dishes) during 12-h RAD21 recovery. (**C**) Box plot of ZF probe distance during RAD21 recovery. For box plots, center lines show the medians; the boxes indicate 25-75%; whiskers extend 1.5 times the interquartile range from the 25th and 75th percentiles. The statistical analysis by Wilcoxon signed-rank test indicated a significant difference between 0 h and 2, 4, 6, and 12 h (*** p <0.001; see Supplementary Table S6).

## Conclusion

In this report, we demonstrated how a pair of ZF probes fused to repeat FLAG and HA epitopes can be coupled with anti-FLAG and anti-HA frankenbodies to mark and track the looping dynamics of two endogenous genomic loci in single living cells. To achieve this, we first created an anti-FLAG frankenbody that complements our recently developed anti-HA frankenbody and that enables the imaging of FLAG-tagged proteins in multiple colors and in diverse live-cell settings. We next designed a pair of synthetic ZF probes with 10× linear HA epitopes (ZF-F) and 10× linear FLAG epitopes (ZF-R) that bind uniquely to two endogenous genomic loci that are close to one another in cellular 3D space. The combination of these two imaging systems allowed us to label and track the relative dynamics of the two target genomic loci in single cells for hours. Unexpectedly large complexes or aggregates appeared to form at target DNA loci by multimerization of ZF probes and frankenbodies. This multimerization is probably mediated by the formation of large complexes of fluorescence proteins under high local concentrations (Filonov et al., 2011; Landgraf et al., 2012; Zacharias et al., 2002). Fortunately, the formation of large complexes containing many ZF probes at target DNA loci not only amplifies fluorescence signal at the site, but also stabilizes DNA binding because each ZF probes within the complex can rapidly rebind DNA following any unbinding event.

The ZF probe-based technology has several advantages. First, we showed our ZF probes can amplify fluorescent signals at target DNA loci at two levels, first at the level of the 10× epitope tag and second at the level of ZF probe multimerization. The resulting ZF probe-frankenbody complexes appear similar to the multimerization of ParB in the ANCHOR system that have been shown to interfere less (Saad et al., 2014). Second, our ZF probes are cell permeable, so we could characterize them biochemically *in vitro* and later load them into living cells in a controlled manner for optimal experiments. Third, our ZF probes can label endogenous loci without the need for time-consuming genetic engineering of cells to be analyzed. While similar systems have been created by amplifying CRISPR-Cas9 signals using the MS2 system (Ma et al., 2018; Qin et al., 2017), the SunTag system (Tanenbaum et al., 2014), or a combination of the two (Hong et al., 2018), our technology is an orthogonal approach for multiplexed imaging in live cells.

While advantageous in several ways, there are a couple of disadvantages of our ZF probes worth pointing out. First, ZF probe design is target sequence limited. In Web tools based on the Barbas set of ZF modules we used in this study (Durai et al., 2005), only 49 out of 64 nucleotide triplet combinations are available. Fortunately, this limitation is not very problematic in our case because genome visualization based on light microscopy does not require base-pair resolution. Thus, unlike genome editing, which requires a precise target cut site, target site selection for live-cell imaging can be within a few-kb region, providing a fair amount of flexibility in the ZF design process. Furthermore, the available triplet combinations have been increasing, and so more flexible designs would be possible (Bhakta & Segal, 2010). Second, our ZF probes tended to aggregate in the cytoplasm of living cells. Fortunately, this did not interfere significantly with our imaging of genomic loci in the cell nucleus. That being said, if a target genomic locus is very close to the nuclear periphery, it may be more difficult to distinguish it from artifactual cytoplasmic aggregates. To circumvent this problem, microscopes with a good sectioning and/or superresolution system, e.g., confocal or light-sheet microscope, will be helpful (Boettiger & Murphy, 2020). Third, the present system uses both transfection of frankenbodies and loading of ZF probes, whose purification is tedious and time consuming. It may be possible to transfect expression vectors of ZF probes together with frankenbodies, but cells that express four proteins with appropriate levels need to be selected. Fourth, it is difficult to demonstrate that the observed foci are the actual targets in living cells. In this study, we used mouse cells lacking a genomic target as a control, and the cohesin-dependent association of two foci support the specific targeting. However, in the future it would be ideal to demonstrate the specificity by individually deleting each target locus.

To demonstrate the potential of our ZF probes, we used them to investigate the role of the cohesin complex in genome organization. This required us to design two unique ZF probes to bind to two endogenous genomic loci that are predicted to be in close contact in 3D cellular space. Consistent with these predictions, when we imaged our ZF probe pair with anti-FLAG and anti-HA frankenbodies in living cells, they were associated with each other in bright nuclear loci in more than half of all cells. Time-lapse imaging further revealed that the two loci became separated after triggered RAD21 degradation. Some separated foci were re-associated by cohesin restoration, as previously demonstrated by Hi-C (Rao et al., 2017). The association rate after cohesin restoration (31.8%) was smaller than that in untreated cells (53.1%), suggesting that cohesin-mediated chromatin association is not deterministic. Thus, our work provides strong evidence that our ZF probes are indeed properly localized and that their relative distance can be used to directly visualize cohesin-mediated chromatin loop dynamics in single, living cells.

In addition, we also showed data implicating the involvement of CTCF and WAPL in maintaining chromatin structure. CTCF can be an anchor and definer for the cohesion border that provides control over looping activities (de Wit et al., 2015; Parelho et al., 2008; Rao et al., 2017). In line with this, our data directly demonstrates CTCF knockdown leads to the loosening of chromatin contacts, but not to the same degree as RAD21 degradation. These data therefore suggest the existence of a CTCF-independent loop (Hansen et al., 2019; Schmidt et al., 2010). The depletion of WAPL also had a mild but significant impact on chromatin contact and double depletion of CTCF and WAPL had an additive effect, which is consistent with a recent study showing that WAPL depletion resulted in cohesin remaining at CTCF sites (Liu et al., 2021).

Having demonstrated how to design and use ZF probes to image the dynamics of non-repetitive genomic loci, our technology can now be adapted to image other endogenous genomic loci of interest. In this study we used published Hi-C data to design a pair of ZF probes to study cohesin-mediated chromatin looping (Sati & Cavalli, 2017; Szabo et al., 2019). Using similar datasets, it is straightforward to design other, similar ZF probes to investigate a wide range of genome dynamics. We therefore anticipate our technology will be of general use to investigate 3D genome dynamics in multiple colors in living cells.

## Acknowledgements

We thank Yuko Sato and Harumi Ueno for cloning anti-FLAG scFv, Takeshi Shimi for instructing the lentivirus expression system, members of Kimura and Stasevich labs for valuable discussion, and the Biomaterials Analysis Division, Open Facility Center, Tokyo Institute of Technology for DNA sequencing.

## Supplementary Information

**Table S1.**
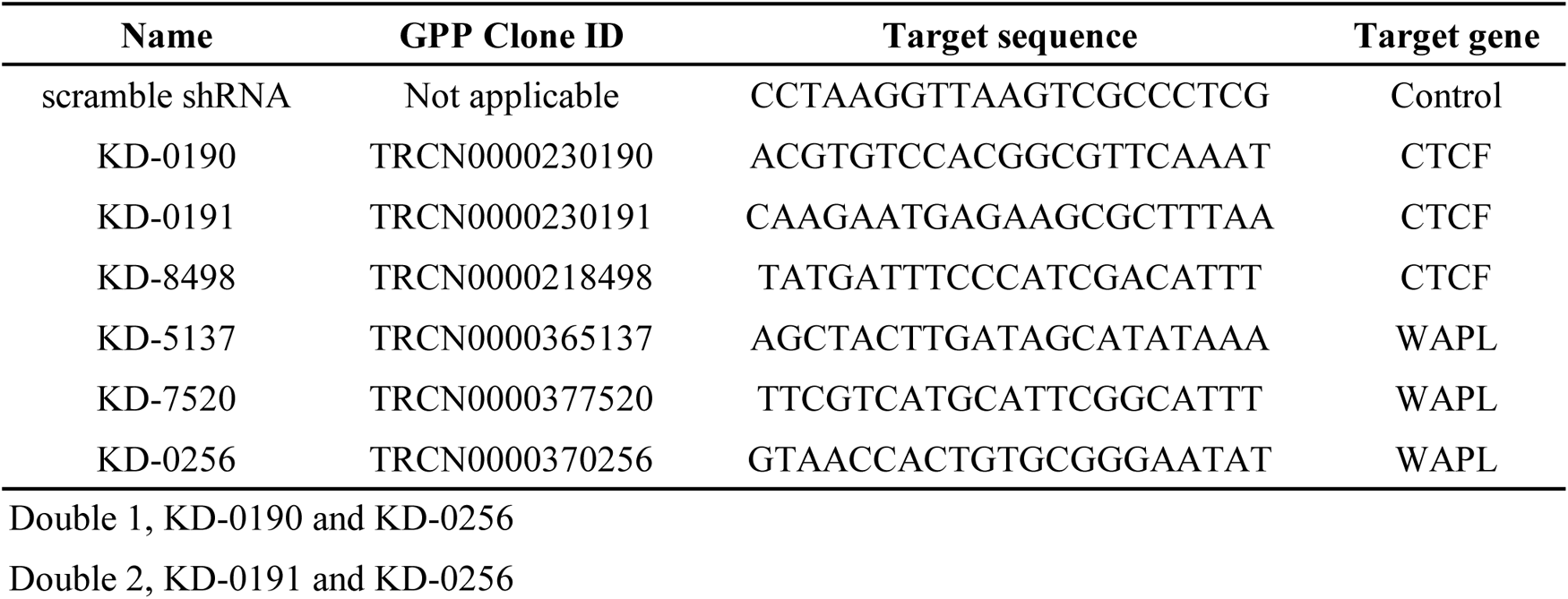
Oligonucleotides for shRNA expression.

**Table S2.**
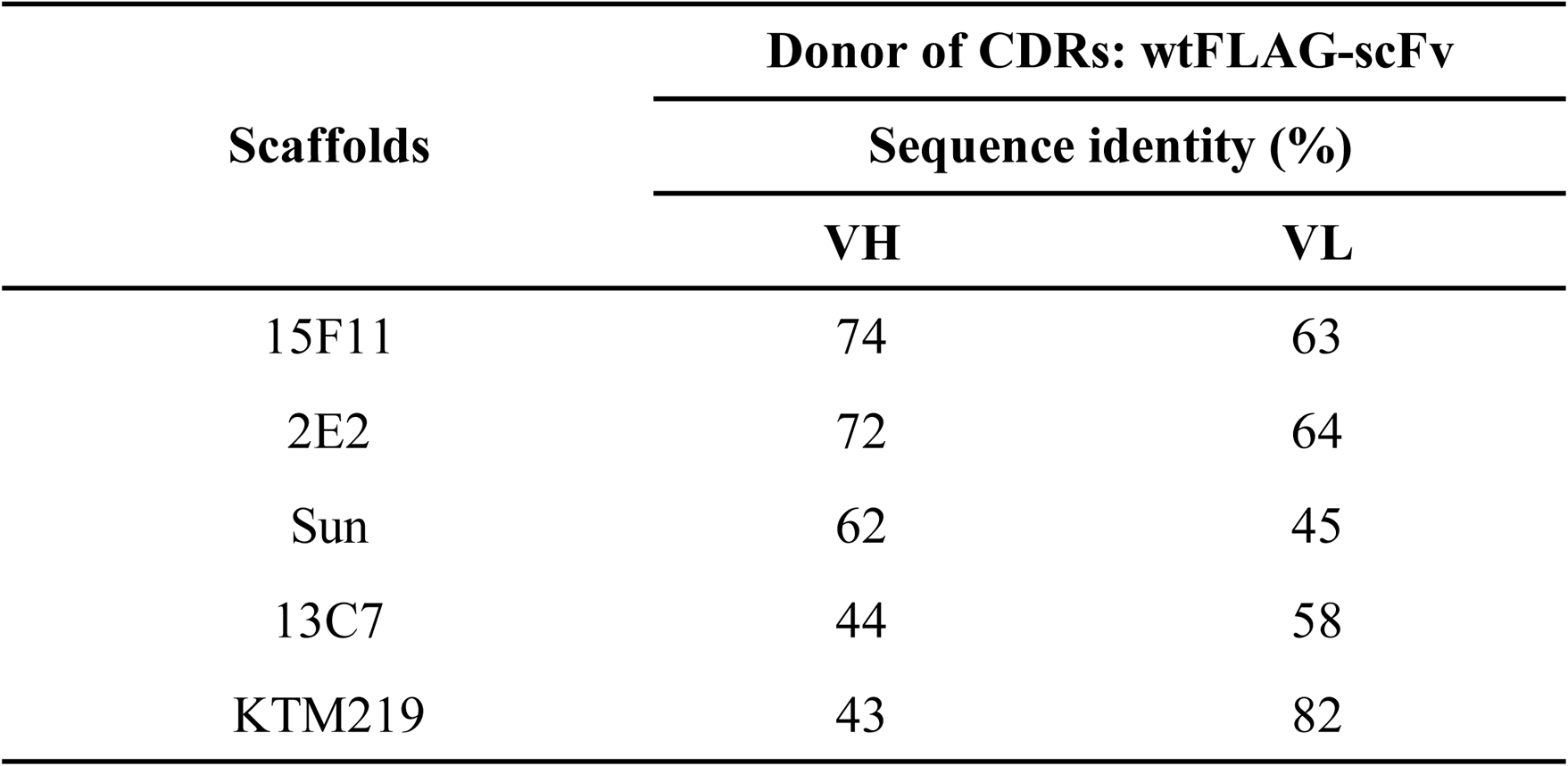
Sequence identity analysis of wtFLAG-scFv with five intracellular scFv scaffolds.

**Table S3.** Descriptive statistics of foci to nucleus intensity ratio.

|  | **mEGFP** | **EGFP** | **mCherry** | **iRFP** |
| --- | --- | --- | --- | --- |
| Mean | 1.44 | 4.28 | 4.89 | 3.53 |
| SD | 0.34 | 2.67 | 2.84 | 2.13 |
| Minimum | 0.98 | 1.22 | 1.48 | 1.03 |
| Maximum | 2.26 | 9.15 | 9.85 | 7.96 |

**Table S4.** Descriptive statistics of fluorescent molecule number per focus.

| **Relative intensity to mean single molecule intensity** |  |
| --- | --- |
| Mean | 138 |
| Median | 104 |
| SD | 123 |
| Minimum | 6.88 |
| Maximum | 499 |

**Table S5.** Statistical analysis of probe distance inHCT116-RAD21-mAID cells.

|  | N | Mean | SD | SE | Wilcoxon signed-rank test (with W/O Auxin) |  |  |
| --- | --- | --- | --- | --- | --- | --- | --- |
|  |  |  |  |  | W | *p* | Vovk-Sellke Maximum *p*-ratio |
| W/O Auxin | 54 | 0.875 | 0.224 | 0.031 | NA<sup>a</sup> | NA | NA |
| W/ Auxin 1 h | 54 | 2.219 | 1.332 | 0.180 | 3.000 | <0.001 | 3.34E+11 |
| W/ Auxin 2 h | 54 | 3.776 | 2.065 | 0.281 | 30.000 | <0.001 | 1.02E+09 |
| W/ Auxin 4 h | 54 | 3.569 | 1.856 | 0.253 | 30.000 | <0.001 | 1.02E+09 |
| W/ Auxin 6 h | 54 | 3.491 | 1.613 | 0.219 | 30.000 | <0.001 | 1.02E+09 |

<sup>a</sup>Not Applicable

**Table S6.** Statistical analysis for the effect of Rad21 recovery on probe distance.

| Time after auxin removal | N | Mean | SD | SE | Wilcoxon signed-rank test (with Recover 0 h) |  |  |
| --- | --- | --- | --- | --- | --- | --- | --- |
|  |  |  |  |  | W | *p* | Vovk-Sellke Maximum *p*-ratio |
| 5 min | 22 | 4.690 | 2.128 | 0.454 | NA<sup>a</sup> | NA | NA |
| 1 h | 22 | 4.301 | 2.180 | 0.465 | 192.000 | 0.033 | 3.28E+00 |
| 2 h | 22 | 3.162 | 1.749 | 0.373 | 253.000 | <0.001 | 5.30E+04 |
| 4 h | 22 | 2.460 | 1.385 | 0.295 | 253.000 | <0.001 | 5.30E+04 |
| 6 h | 22 | 2.461 | 1.456 | 0.310 | 253.000 | <0.001 | 5.30E+04 |
| 12 h | 17 | 2.173 | 1.275 | 0.319 | 136.000 | <0.001 | 1.16E+03 |

<sup>a</sup>Not Applicable

**Fig. S1.**
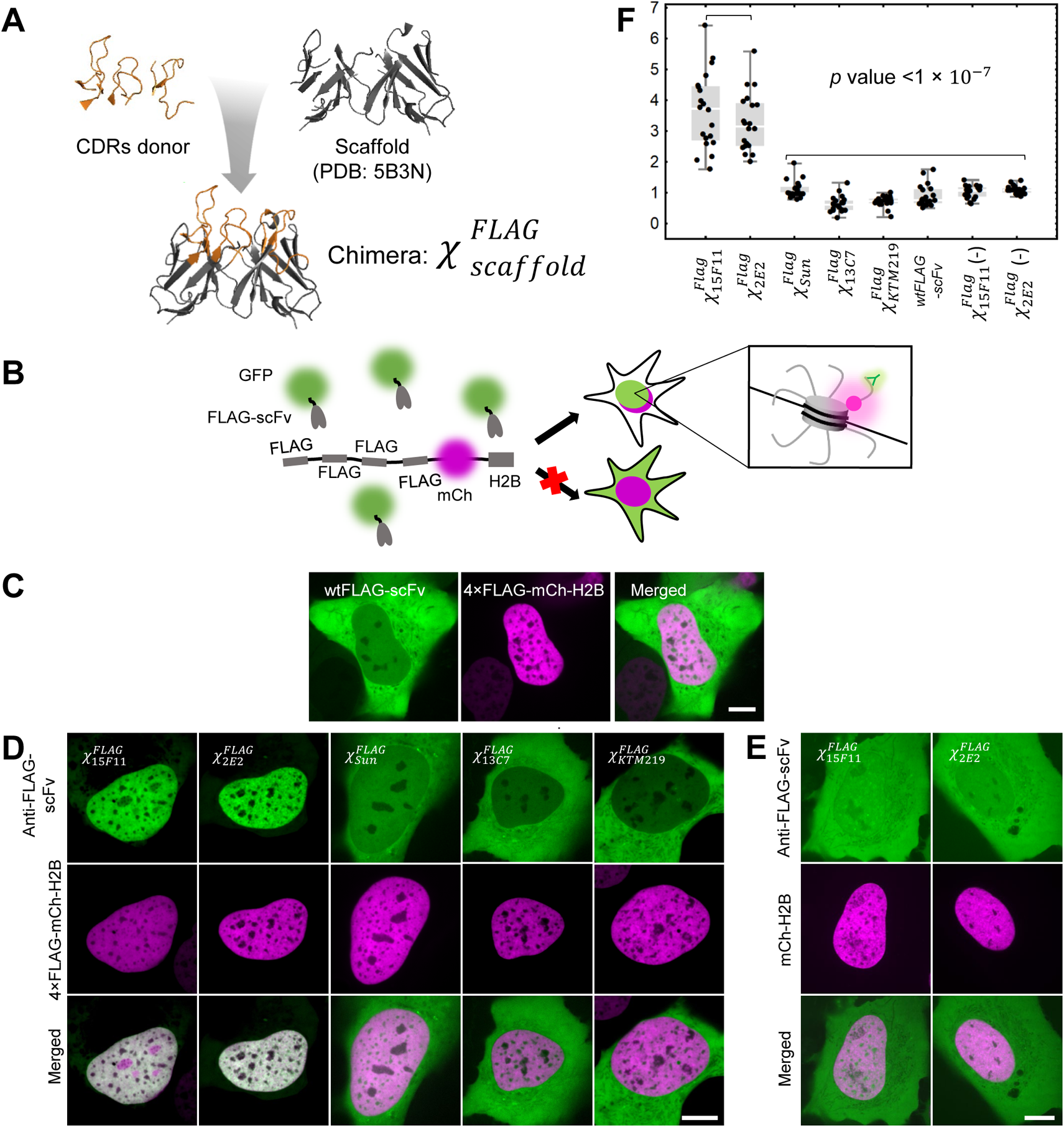
Design strategy and initial screening of anti-FLAG frankenbodies. (**A**) A cartoon schematic showing how to design a chimeric anti-FLAG scFv using wtFLAG-scFv CDRs and stable scFv scaffolds. (**B**) A cartoon showing how to screen the five chimeric anti-FLAG scFvs in living U2OS cells. (**C**)A representative cell showing the respective localization of the wildtype anti-FLAG-scFv in living U2OS cells co-expressing FLAG-tagged histone H2B (wtFLAG-scFv, green; 4×FLAG-mCh-H2B, magenta). (**D**) Initial screening results showing the respective localization of the five chimeric anti-FLAG scFvs in living U2OS cells co-expressing FLAG-tagged histone H2B (chimeric anti-FLAG scFv, green; 4×FLAG-mCh-H2B, magenta). (**E**) Control results showing the respective localization of anti-FLAG frankenbodies in living cells lacking FLAG-tagged histone H2B (chimeric anti-FLAG scFv, green; mCh-H2B, magenta). (**F**) Nuclear to cytoplasmic fluorescent intensity ratio (Nuc/Cyt) plot of each chimeric anti-FLAG scFv and wtFLAG-scFv for all cells imaged as in (**C**), (**D**) and (**E**). Mann-Whitney test. All images are representative cell images from one independent experiment. Scale bars: 10µm. Source data are provided as a Source Data file. For box plots, center lines show the medians; the boxes indicate 25-75%; whiskers extend 1.5 times the interquartile range from the 25th and 75th percentiles; and data points are plotted.

**Fig. S2.**
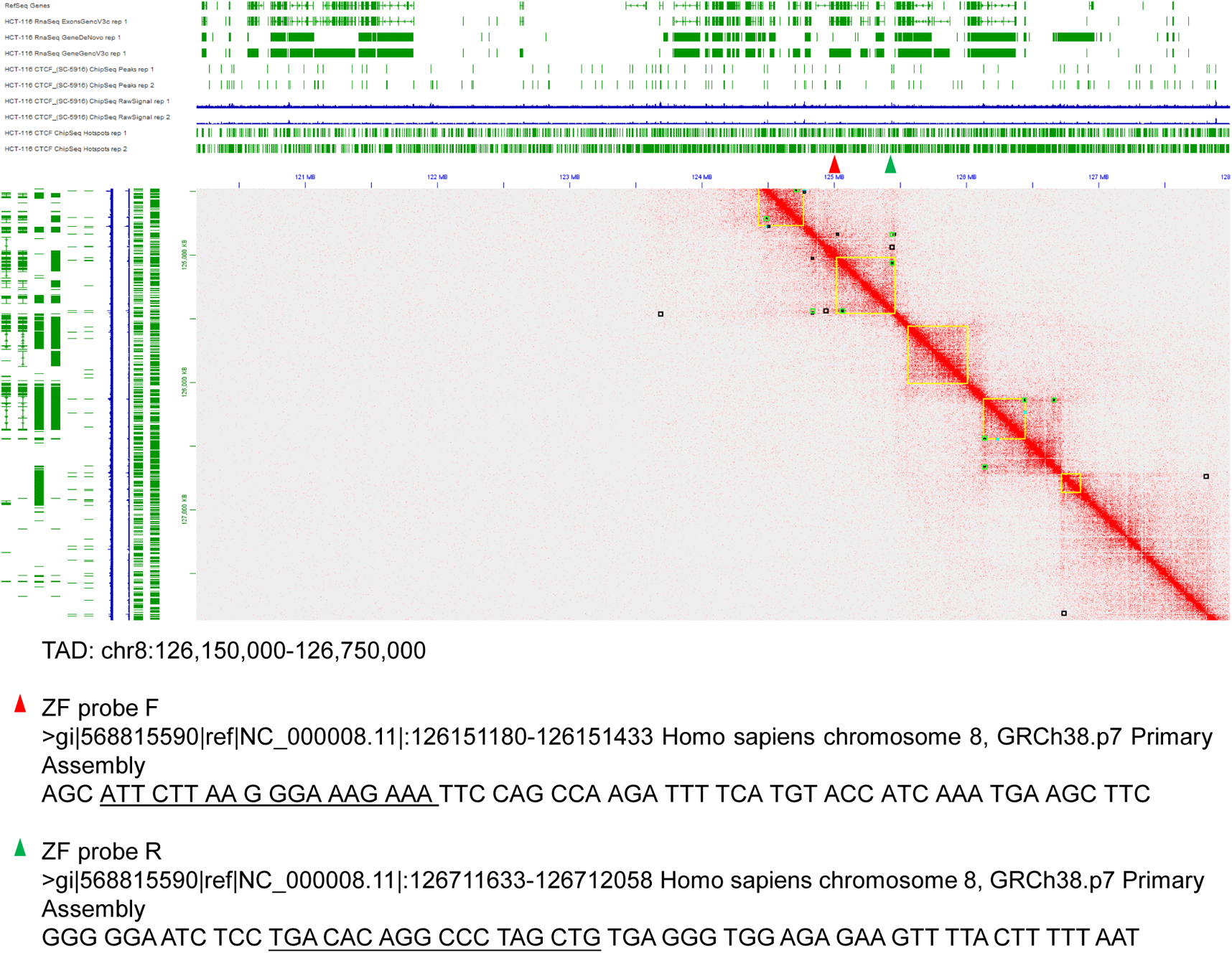
ZF probe target sites. Chromatin contact sites identified by multiple HiC algorithms, HiC pro (Black), Juicebox(Green), and data from Li et al, 2012 (Yellow) is aligned with ENCODEHCT116 CTCF ChipSeq data (ENCSR000BSE) and RNASeq (ENCSR000CWM) data. ZF probes that bind to the upstream and downstream sequence (underlined) are named F and R, respectively.

**Fig. S3.**
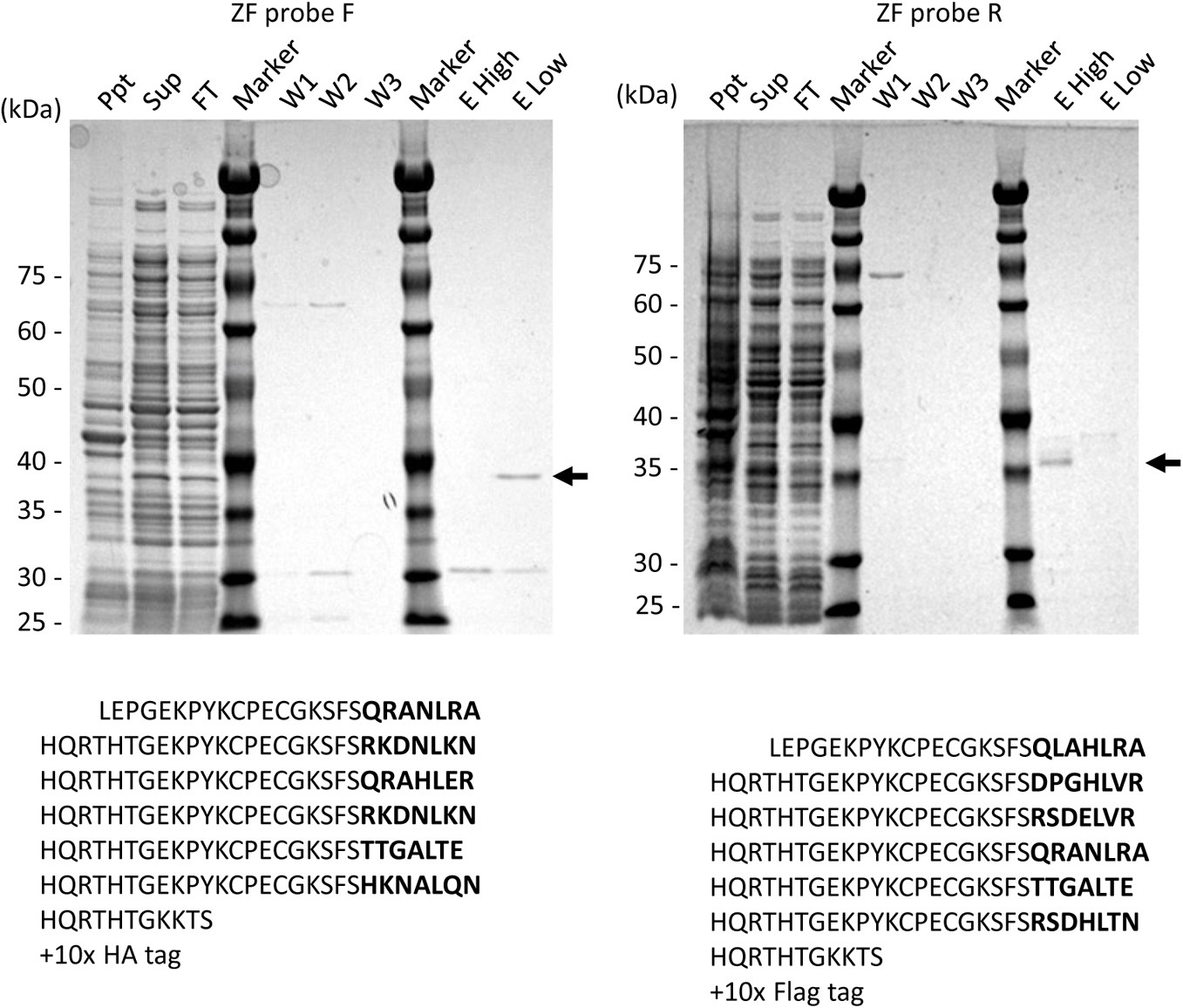
Expression and purification of ZF probes. Lysates of *E. coli* expressing ZF probes were centrifuged to remove the insoluble precipitate (Ppt). The supernatant (Sup) was applied to an Ni column and the flow-through (FL), Wash 1 (W1), Wash 2 (W2), Wash 3 (W3) fractions were collected. Bound proteins were then eluted and the elution peak 1(E high) and peak 2 (E low) were collected. All fractions were analyzed by SDS-PAGE and Coomassie Blue staining. The sizes of marker proteins (Marker) are indicated on the left. The positions of ZF probes are indicated by arrows on the right. Amino acid sequence of ZF proteins are shown with the variable regions that determine the DNA-binding specificity indicated in bold.

**Fig. S4.**
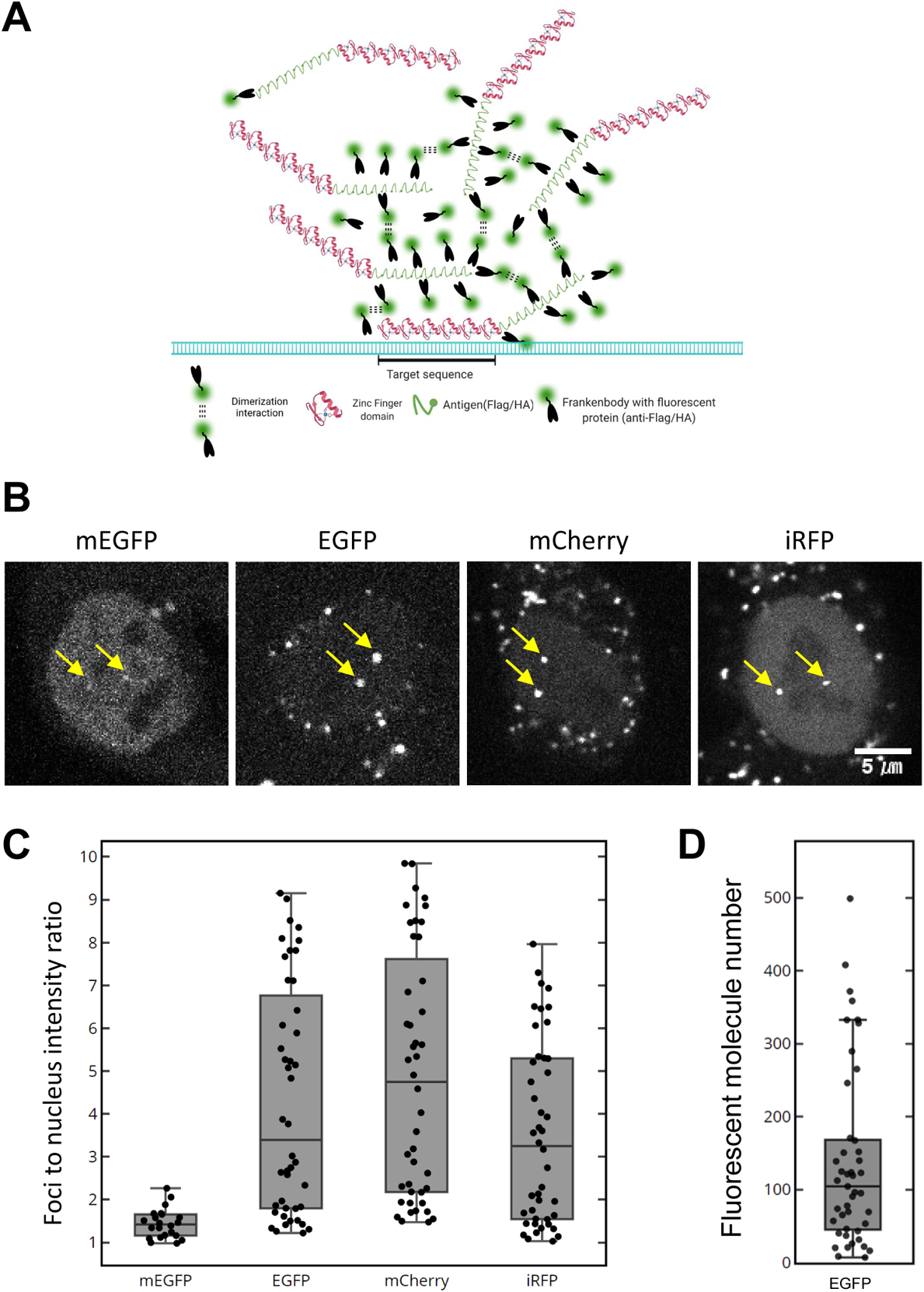
Multimer formation and ZF probes with frankenbodies. (A) A schematic illustration for a model that ZF probes and frankenbodies make a big complex through dimerization of fluorescent proteins. Such a complex contains multiple DNA binding domains and fluorescent proteins. (B and C) Wild-type HCT-116 cells were transfected with expression vectors for anti-HA frankenbody tagged with a fluorescent protein (mEGFP, EGFP, mCherry, and iRFP) and then loaded with ZF-F probe that harbor 10x HA. (B) Representative confocal images of living HCT-116 cells transfected with anti-HA frankenbody. Nuclear foci are indicated by yellow arrows. (C) Intensity ratios of foci to nuclear background of anti-HA frankenbodies tagged with a different fluorescent protein (N_mEGFP_ = 23, N_EGFP_ = 44, N_mCherry_ = 45 and N_iRFP_ = 44; each from 2 dishes). See Supplementary Table S3. (D) Relative intensity of foci to that of single molecule fluorescence. The center line shows the median; the box indicates 25-75%; whiskers extend 1.5 times the interquartile range from the 25th and 75th percentiles; and data points are plotted. The average is 138 (N = 45). See Supplementary Table S4.

**Fig. S5.**
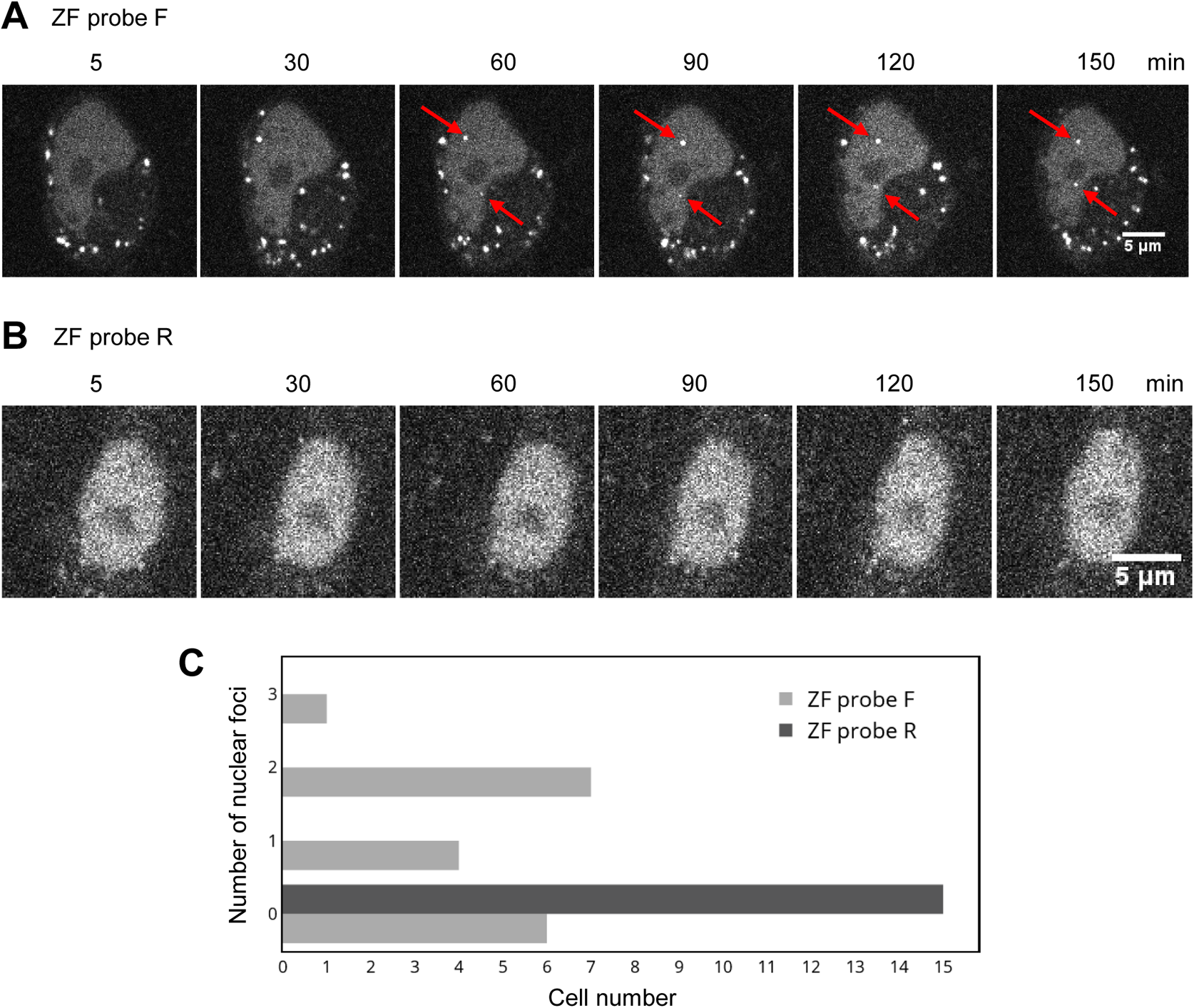
ZF probes in mouse cells. Mouse A9 cells were transfected with anti-HA mCherry or anti-FLAG iRFP frankenbody. After administrating ZF probes, time-lapse confocal images were collected. In the mouse genome, the target sequence of ZF probe F, but not R, is present. (A) ZF probe F with anti-HA mCherry-frankenbody. Nuclear foci are indicated by arrows. (B) ZF probe R with anti-FLAG iRFP-frankenbody. No nuclear foci were observed. (C) Number of nuclear foci per cell (n=18 and 15 for ZF probe F and R, respectively).

**Fig. S6.**
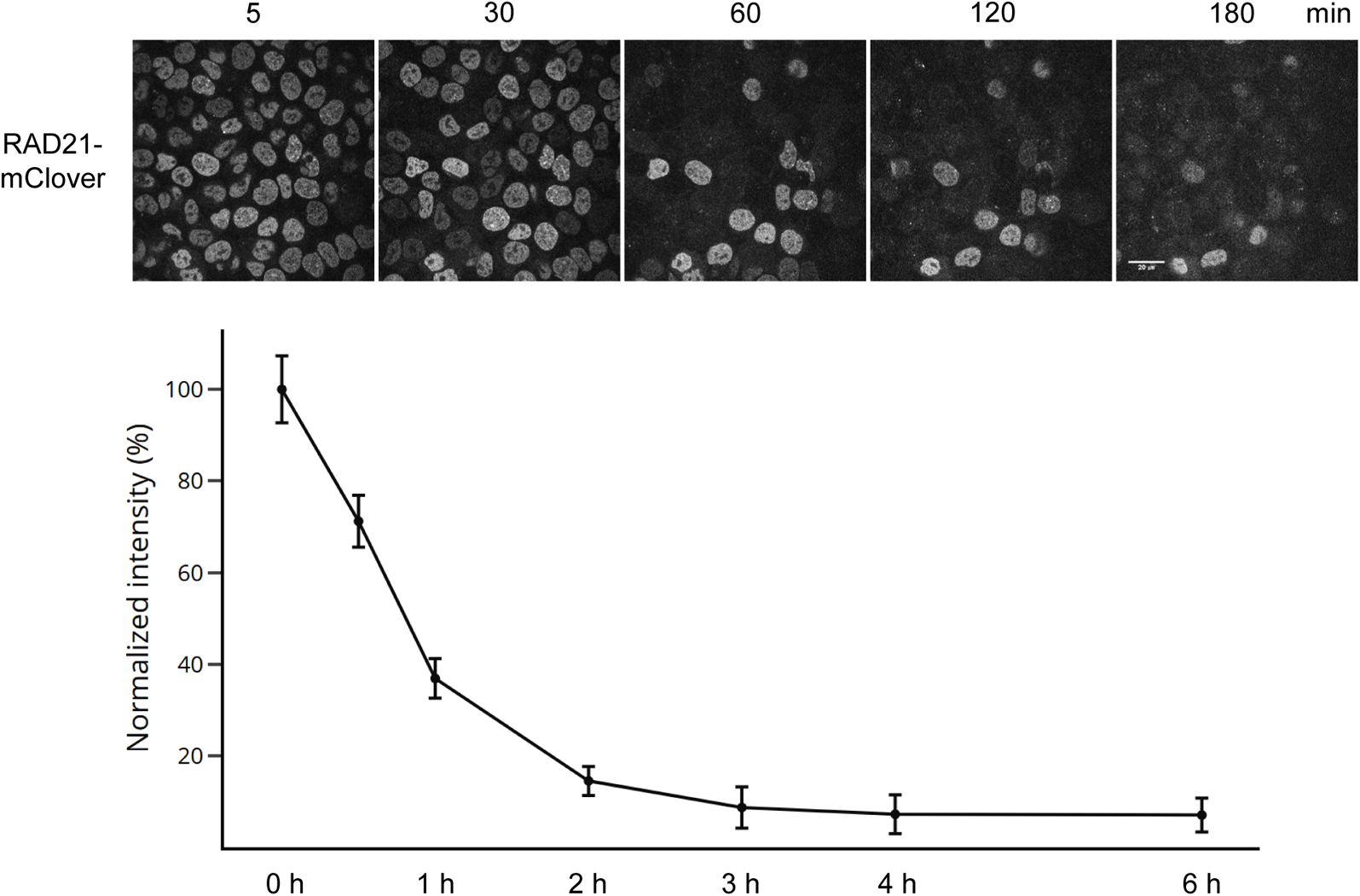
Degradation of RAD21-mClover by auxin treatment. After the addition of auxin in the medium, time-lapse confocal sections of HCT116-RAD21-mAID-mClover cells were acquired. (top) Representative images of RAD21-mClover. (bottom) Normalized intensity of RAD21-mClover to that of time point 0 h (average ± SD; n=1024 pixel-points; from duplicate experiments).

**Fig. S7.**
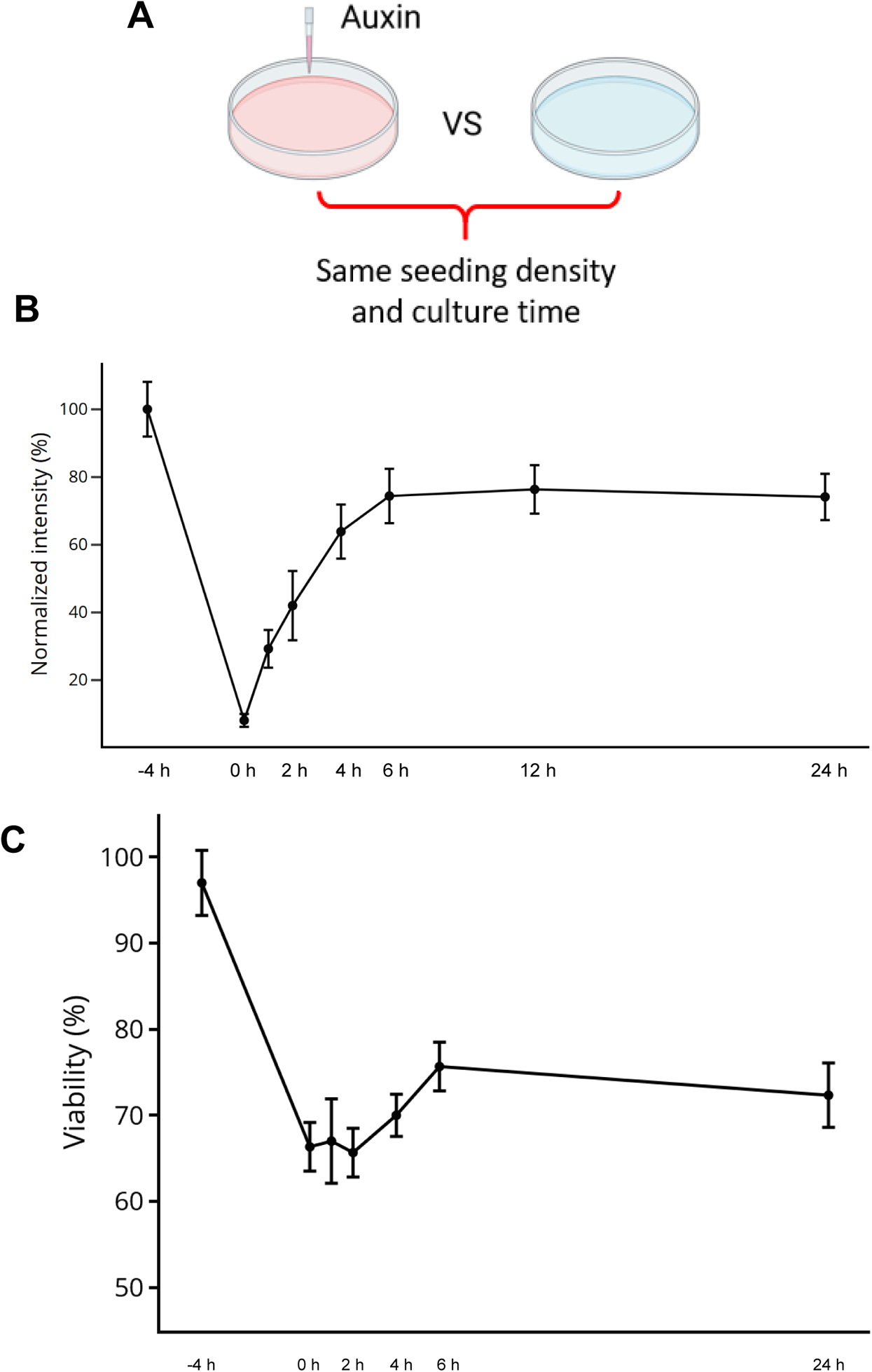
Recovery of HCT116-RAD21-mAID-mClover cells after removal of auxin. HCT116-RAD21-mAID-mClover cells were transfected with anti-FLAG and anti-HA frankenbody expression vectors, loaded with ZF probes, and treated with auxin for 4 h to degrade RAD21. After washing away auxin, cells were set on to a confocal microscope and further incubated with auxin-free medium. (**A**) A schematic illustration for how to seeding negative control (NC) sample for both RAD21-mClover-GFP intensity and cell viability. (**B**) Normalized RAD21-mClover-GFP intensity (average ± SD; n=107 cells). (**C**) Cell viability. Cell numbers were counted after the auxin removal (average ± SD; n=3 independent experiments).

## References

Anton, T., Bultmann, S., Leonhardt, H., & Markaki, Y. (2014). Visualization of specific DNA sequences in living mouse embryonic stem cells with a programmable fluorescent CRISPR/Cas system. Nucleus 5, 163–172.

Banigan, E.J., Berg, A.A. van den, Brandão, H.B., Marko, J.F., & Mirny, L.A. (2020). Chromosome organization by one-sided and two-sided loop extrusion. Elife 9, e53558.

Barrow, J.J., Masannat, J., & Bungert, J. (2012). Neutralizing the function of a β-globin–associated cis-regulatory DNA element using an artificial zinc finger DNA-binding domain. Proc National Acad Sci 109, 17948–17953.

Baumann, K. (2020). Human cohesin extrudes interphase DNA to make loops. Nat Rev Mol Cell Bio 21, 3–3.

Bhakta, M.S., & Segal, D.J. (2010). Engineered Zinc Finger Proteins, Methods and Protocols. Methods Mol Biology 649, 3–30.

Bickmore, W.A. (2012). The Spatial Organization of the Human Genome. Annu Rev Genom Hum G 14, 67–84.

Boettiger, A., & Murphy, S. (2020). Advances in Chromatin Imaging at Kilobase-Scale Resolution. Trends Genet 36, 273–287.

Brandão, H.B., Gabriele, M., & Hansen, A.S. (2021). Tracking and interpreting long-range chromatin interactions with super-resolution live-cell imaging. Curr Opin Cell Biol 70, 18–26.

Cencic, R., Miura, H., Malina, A., Robert, F., Ethier, S., Schmeing, T.M., Dostie, J., & Pelletier, J. (2014). Protospacer Adjacent Motif (PAM)-Distal Sequences Engage CRISPR Cas9 DNA Target Cleavage. Plos One 9, e109213.

Chen, B., Gilbert, L.A., Cimini, B.A., Schnitzbauer, J., Zhang, W., Li, G.-W., Park, J., Blackburn, E.H., Weissman, J.S., Qi, L.S., & Huang, B. (2013). Dynamic Imaging of Genomic Loci in Living Human Cells by an Optimized CRISPR/Cas System. Cell 155, 1479–1491.

Choo, Y., Sánchez-García, I., & Klug, A. (1994). In vivo repression by a site-specific DNA-binding protein designed against an oncogenic sequence. Nature 372, 642–645.

Cialek, C.A., Galindo, G., Koch, A.L., Saxton, M.N., & Stasevich, T.J. (2021). Bead Loading Proteins and Nucleic Acids into Adherent Human Cells. J Vis Exp 172, doi: 10.3791/62559.

Colca, J.R., McDonald, W.G., Waldon, D.J., Leone, J.W., Lull, J.M., Bannow, C.A., Lund, E.T. & Mathews, W.R. (2004). Identification of a novel mitochondrial protein (“mitoNEET”) cross-linked specifically by a thiazolidinedione photoprobe. Am J Physiol Endocrinol Metab 286, E252–E260.

Dekker, J., Rippe, K., Dekker, M., & Kleckner, N. (2002). Capturing Chromosome Conformation. Science 295, 1306–1311.

de Wit, E., Vos, E.S.M., Holwerda, S.J.B., Valdes-Quezada, C., Verstegen, M.J.A.M., Teunissen, H., Splinter, E., Wijchers, P.J., Krijger, P.H.L., & de Laat, W. (2015). CTCF Binding Polarity Determines Chromatin Looping. Mol Cell 60, 676–684.

Dillon, N., Trimborn, T., Strouboulis, J., Fraser, P., & Grosveld, F. (1997). The Effect of Distance on Long-Range Chromatin Interactions. Mol Cell 1, 131–139.

Dixon, J.R., Selvaraj, S., Yue, F., Kim, A., Li, Y., Shen, Y., Hu, M., Liu, J.S., & Ren, B. (2012). Topological domains in mammalian genomes identified by analysis of chromatin interactions. Nature 485, 376–380.

Doi, G., Okada, S., Yasukawa, T., Sugiyama, Y., Bala, S., Miyazaki, S., Kang, D., & Ito, T. (2021). Catalytically inactive Cas9 impairs DNA replication fork progression to induce focal genomic instability. Nucleic Acids Res 49, gkaa1241.

Durai, S., Mani, M., Kandavelou, K., Wu, J., Porteus, M.H., & Chandrasegaran, S. (2005). Zinc finger nucleases: custom-designed molecular scissors for genome engineering of plant and mammalian cells. Nucleic Acids Res 33, 5978–5990.

Ewert, S., Honegger, A., & Plückthun, A. (2004). Stability improvement of antibodies for extracellular and intracellular applications: CDR grafting to stable frameworks and structure-based framework engineering. Methods 34, 184–199.

Filonov, G.S., Piatkevich, K.D., Ting, L.-M., Zhang, J., Kim, K., & Verkhusha, V.V. (2011). Bright and stable near-infrared fluorescent protein for in vivo imaging. Nat Biotechnol 29, 757–761.

Finn, E.H., Pegoraro, G., Brandão, H.B., Valton, A.-L., Oomen, M.E., Dekker, J., Mirny, L., & Misteli, T. (2019). Extensive Heterogeneity and Intrinsic Variation in Spatial Genome Organization. Cell 176, 1502–1515.e10.

Gaj, T., & Liu, J. (2015). Direct Protein Delivery to Mammalian Cells Using Cell-permeable Cys_2_-His_2_ Zinc-finger Domains. J Vis Exp 97, doi: 10.3791/52814.

Gaj, T., Liu, J., Anderson, K.E., Sirk, S.J., & Barbas, C.F. (2014). Protein Delivery Using Cys2–His2 Zinc-Finger Domains. Acs Chem Biol 9, 1662–1667.

Garcia-Bloj, B., Moses, C., Sgro, A., Plani-Lam, J., Arooj, M., Duffy, C., Thiruvengadam, S., Sorolla, A., Rashwan, R., Mancera, R.L., Leisewitz, A., Swift-Scanlan, T., Corvalan, A.H., & Blancafort, P. (2016). Waking up dormant tumor suppressor genes with zinc fingers, TALEs and the CRISPR/dCas9 system. Oncotarget 7, 60535–60554.

Garriga-Canut, M., Agustín-Pavón, C., Herrmann, F., Sánchez, A., Dierssen, M., Fillat, C., & Isalan, M. (2012). Synthetic zinc finger repressors reduce mutant huntingtin expression in the brain of R6/2 mice. Proc National Acad Sci 109, E3136–E3145.

Gerlich, D., Koch, B., Dupeux, F., Peters, J.-M., & Ellenberg, J. (2006). Live-Cell Imaging Reveals a Stable Cohesin-Chromatin Interaction after but Not before DNA Replication. Curr Biol 16, 1571–1578.

Gersbach, C.A., Gaj, T., & Barbas, C.F. (2014). Synthetic Zinc Finger Proteins: The Advent of Targeted Gene Regulation and Genome Modification Technologies. Accounts Chem Res 47, 2309–2318.

Gizzi, A.M.C., Cattoni, D.I., Fiche, J.-B., Espinola, S.M., Gurgo, J., Messina, O., Houbron, C., Ogiyama, Y., Papadopoulos, G.L., Cavalli, G., Lagha, M., & Nollmann, M. (2019). Microscopy-Based Chromosome Conformation Capture Enables Simultaneous Visualization of Genome Organization and Transcription in Intact Organisms. Mol Cell 74, 212–222.e5.

Grimm, J.B., English, B.P., Chen, J., Slaughter, J.P., Zhang, Z., Revyakin, A., Patel, R., Macklin, J.J., Normanno, D., Singer, R.H., Lionnet, T., & Lavis, L.D. (2015). A general method to improve fluorophores for live-cell and single-molecule microscopy. Nat Methods 12, 244–250.

Hansen, A.S., Cattoglio, C., Darzacq, X., & Tjian, R. (2017). Recent evidence that TADs and chromatin loops are dynamic structures. Nucleus 9, 1–18.

Hansen, A.S., Hsieh, T.-H.S., Cattoglio, C., Pustova, I., Saldaña-Meyer, R., Reinberg, D., Darzacq, X., & Tjian, R. (2019). Distinct Classes of Chromatin Loops Revealed by Deletion of an RNA-Binding Region in CTCF. Mol Cell 76, 395–411.e13.

Hong, Y., Lu, G., Duan, J., Liu, W., & Zhang, Y. (2018). Comparison and optimization of CRISPR/dCas9/gRNA genome-labeling systems for live cell imaging. Genome Biol 19, 39.

Imanishi, M., Hori, Y., Nagaoka, M., & Sugiura, Y. (2000). DNA-Bending Finger: Artificial Design of 6-Zinc Finger Peptides with Polyglycine Linker and Induction of DNA Bending †. Biochemistry-Us 39, 4383–4390.

Isalan, M., Choo, Y., & Klug, A. (1997). Synergy between adjacent zinc fingers in sequence-specific DNA recognition. Proc National Acad Sci 94, 5617–5621.

Jiang, F., Zhou, K., Ma, L., Gressel, S., & Doudna, J.A. (2015). A Cas9–guide RNA complex preorganized for target DNA recognition. Science 348, 1477–1481.

Joglekar, A.P., Salmon, E. D. & Bloom, K. S. (2008). Counting kinetochore protein numbers in budding yeast using genetically encoded fluorescent proteins. Methods Cell Biol 85, 127–151.

Joyce, E.F., Williams, B.R., Xie, T., & Wu, C. -ting (2012). Identification of Genes That Promote or Antagonize Somatic Homolog Pairing Using a High-Throughput FISH–Based Screen. Plos Genet 8, e1002667.

Koulouras, G., Panagopoulos, A., Rapsomaniki, M.A., Giakoumakis, N.N., Taraviras, S., & Lygerou, Z. (2018). EasyFRAP-web: a web-based tool for the analysis of fluorescence recovery after photobleaching data. Nucleic Acids Res 46, gky508-.

Kueng, S., Hegemann, B., Peters, B.H., Lipp, J.J., Schleiffer, A., Mechtler, K., & Peters, J.-M. (2006). Wapl Controls the Dynamic Association of Cohesin with Chromatin. Cell 127, 955–967.

Landgraf, D., Okumus, B., Chien, P., Baker, T.A., & Paulsson, J. (2012). Segregation of molecules at cell division reveals native protein localization. Nat Methods 9, 480–482.

Lee, H.B., Sundberg, B.N., Sigafoos, A.N., & Clark, K.J. (2016). Genome Engineering with TALE and CRISPR Systems in Neuroscience. Frontiers Genetics 7, 47.

Li, G., Ruan, X., Auerbach, R.K., et al. (2012). Extensive Promoter-Centered Chromatin Interactions Provide a Topological Basis for Transcription Regulation. Cell 148, 84–98.

Lieberman-Aiden, E., Berkum, N.L. van, Williams, L., et al. (2009). Comprehensive Mapping of Long-Range Interactions Reveals Folding Principles of the Human Genome. Science 326, 289–293.

Lim, W.M., Ito, Y., Sakata-Sogawa, K. & Tokunaga, M. (2018). CLIP-170 is essential for MTOC repositioning during T cell activation by regulating dynein localisation on the cell surface. Sci Rep 8, 17447.

Lindhout, B.I., Fransz, P., Tessadori, F., Meckel, T., Hooykaas, P.J.J., & Zaal, B.J. van der (2007). Live cell imaging of repetitive DNA sequences via GFP-tagged polydactyl zinc finger proteins. Nucleic Acids Res 35, e107–e107.

Liu, N.Q., Maresca, M., Brand, T. van den, Braccioli, L., Schijns, M.M.G.A., Teunissen, H., Bruneau, B.G., Nora, E.P., & Wit, E. de (2021). WAPL maintains a cohesin loading cycle to preserve cell-type-specific distal gene regulation. Nat Genet 53, 100–109.

Ma, H., Tu, L.-C., Naseri, A., Chung, Y.-C., Grunwald, D., Zhang, S., & Pederson, T. (2018). CRISPR-Sirius: RNA scaffolds for signal amplification in genome imaging. Nat Methods 15, 928–931.

Mandell, J.G., & Barbas, C.F. (2006). Zinc Finger Tools: custom DNA-binding domains for transcription factors and nucleases. Nucleic Acids Res 34, W516–W523.

Michaelis, C., Ciosk, R., & Nasmyth, K. (1997). Cohesins: Chromosomal Proteins that Prevent Premature Separation of Sister Chromatids. Cell 91, 35–45.

Mino, T., Mori, T., Aoyama, Y., & Sera, T. (2008). Cell-permeable artificial zinc-finger proteins as potent antiviral drugs for human papillomaviruses. Arch Virol 153, 1291.

Miyanari, Y., Ziegler-Birling, C., & Torres-Padilla, M.-E. (2013). Live visualization of chromatin dynamics with fluorescent TALEs. Nat Struct Mol Biol 20, 1321–1324.

Morisaki, T., Lyon, K., DeLuca, K.F., DeLuca, J.G., English, B.P., Zhang, Z., Lavis, L.D., Grimm, J.B., Viswanathan, S., Looger, L.L., Lionnet, T., & Stasevich, T.J. (2016). Real-time quantification of single RNA translation dynamics in living cells. Science 352, 1425–1429.

Natsume, T., Kiyomitsu, T., Saga, Y., & Kanemaki, M.T. (2016). Rapid Protein Depletion in Human Cells by Auxin-Inducible Degron Tagging with Short Homology Donors. Cell Reports 15, 210–218.

Nora, E.P., Lajoie, B.R., Schulz, E.G., et al. (2012). Spatial partitioning of the regulatory landscape of the X-inactivation centre. Nature 485, 381–385.

Parelho, V., Hadjur, S., Spivakov, M., et al. (2008). Cohesins Functionally Associate with CTCF on Mammalian Chromosome Arms. Cell 132, 422–433.

Perez, E.E., Wang, J., Miller, J.C., et al. (2008). Establishment of HIV-1 resistance in CD4+ T cells by genome editing using zinc-finger nucleases. Nat Biotechnol 26, 808–816.

Pombo, A., & Dillon, N. (2015). Three-dimensional genome architecture: players and mechanisms. Nat Rev Mol Cell Bio 16, 245–257.

Qi, L.S., Larson, M.H., Gilbert, L.A., Doudna, J.A., Weissman, J.S., Arkin, A.P., & Lim, W.A. (2013). Repurposing CRISPR as an RNA-Guided Platform for Sequence-Specific Control of Gene Expression. Cell 152, 1173–1183.

Qin, P., Parlak, M., Kuscu, C., Bandaria, J., Mir, M., Szlachta, K., Singh, R., Darzacq, X., Yildiz, A., & Adli, M. (2017). Live cell imaging of low- and non-repetitive chromosome loci using CRISPR-Cas9. Nat Commun 8, 14725.

Ran, F.A., Hsu, P.D., Wright, J., Agarwala, V., Scott, D.A., & Zhang, F. (2013). Genome engineering using the CRISPR-Cas9 system. Nat Protoc 8, 2281–2308.

Rao, S.S.P., Huntley, M.H., Durand, N.C., Stamenova, E.K., Bochkov, I.D., Robinson, J.T., Sanborn, A.L., Machol, I., Omer, A.D., Lander, E.S., & Aiden, E.L. (2014). A 3D Map of the Human Genome at Kilobase Resolution Reveals Principles of Chromatin Looping. Cell 159, 1665–1680.

Rao, S.S.P., Huang, S.-C., Hilaire, B.G.S., et al. (2017). Cohesin Loss Eliminates All Loop Domains. Cell 171, 305–320.e24.

Rinaldi, F.C., Doyle, L.A., Stoddard, B.L., & Bogdanove, A.J. (2017). The effect of increasing numbers of repeats on TAL effector DNA binding specificity. Nucleic Acids Res 45, gkx342-.

Robinett, C.C., Straight, A., Li, G., Willhelm, C., Sudlow, G., Murray, A., & Belmont, A.S. (1996). In vivo localization of DNA sequences and visualization of large-scale chromatin organization using lac operator/repressor recognition. J Cell Biology 135, 1685–1700.

Robinson, J.T., Turner, D., Durand, N.C., Thorvaldsdóttir, H., Mesirov, J.P., & Aiden, E.L. (2018). Juicebox.js Provides a Cloud-Based Visualization System for Hi-C Data. Cell Syst 6, 256–258.e1.

Rueden, C.T., Schindelin, J., Hiner, M.C., DeZonia, B.E., Walter, A.E., Arena, E.T., & Eliceiri, J. W. (2017). ImageJ2: ImageJ for the next generation of scientific image data. Bmc Bioinformatics 18, 529.

Saad, H., Gallardo, F., Dalvai, M., Tanguy-le-Gac, N., Lane, D., & Bystricky, K. (2014). DNA Dynamics during Early Double-Strand Break Processing Revealed by Non-Intrusive Imaging of Living Cells. Plos Genet 10, e1004187.

Sati, S., & Cavalli, G. (2017). Chromosome conformation capture technologies and their impact in understanding genome function. Chromosoma 126, 33–44.

Sato, Y., Mukai, M., Ueda, J., et al. (2013). Genetically encoded system to track histone modification in vivo. Sci Rep-Uk 3, 2436.

Sato, Y., Kujirai, T., Arai, R., Asakawa, H., Ohtsuki, C., Horikoshi, N., Yamagata, K., Ueda, J., Nagase, T., Haraguchi, T., Hiraoka, Y., Kimura, A., Kurumizaka, H., & Kimura, H. (2016). A Genetically Encoded Probe for Live-Cell Imaging of H4K20 Monomethylation. J Mol Biol 428, 3885–3902.

Schmidt, D., Schwalie, P.C., Ross-Innes, C.S., Hurtado, A., Brown, G.D., Carroll, J.S., Flicek, P., & Odom, D.T. (2010). A CTCF-independent role for cohesin in tissue-specific transcription. Genome Res 20, 578–588.

Sera, T. (2010). Engineered Zinc Finger Proteins, Methods and Protocols. Methods Mol Biology 649, 91–96.

Servant, N., Varoquaux, N., Lajoie, B.R., Viara, E., Chen, C.-J., Vert, J.-P., Heard, E., Dekker, J., & Barillot, E. (2015). HiC-Pro: an optimized and flexible pipeline for Hi-C data processing. Genome Biol 16, 259.

Shaban, H.A., Barth, R., & Bystricky, K. (2020). Navigating the crowd: visualizing coordination between genome dynamics, structure, and transcription. Genome Biol 21, 278.

Szabo, Q., Bantignies, F., & Cavalli, G. (2019). Principles of genome folding into topologically associating domains. Sci Adv 5, eaaw1668.

Tanenbaum, M.E., Gilbert, L.A., Qi, L.S., Weissman, J.S., & Vale, R.D. (2014). A Protein-Tagging System for Signal Amplification in Gene Expression and Fluorescence Imaging. Cell 159, 635–646.

Tebas, P., Stein, D., Tang, W.W., et al. (2014). Gene Editing of CCR5 in Autologous CD4 T Cells of Persons Infected with HIV. New Engl J Medicine 370, 901–910.

Tedeschi, A., Wutz, G., Huet, S., et al. (2013). Wapl is an essential regulator of chromatin structure and chromosome segregation. Nature 501, 564–568.

Tokunaga, M., Imamoto, N., & Sakata-Sogawa, K. (2008). Highly inclined thin illumination enables clear single-molecule imaging in cells. Nat Methods 5, 159–161.

Waldo, G.S., Standish, B.M., Berendzen, J., & Terwilliger, T.C. (1999). Rapid protein-folding assay using green fluorescent protein. Nat Biotechnol 17, 691–695.

Wijgerde, M., Grosveld, F., & Fraser, P. (1995). Transcription complex stability and chromatin dynamics in vivo. Nature 377, 209–213.

Wilen, C.B., Wang, J., Tilton, J.C., et al. (2011). Engineering HIV-Resistant Human CD4+ T Cells with CXCR4-Specific Zinc-Finger Nucleases. Plos Pathog 7, e1002020.

Woglar, A., Yamaya, K., Roelens, B., Boettiger, A., Köhler, S., & Villeneuve, A.M. (2020). Quantitative cytogenetics reveals molecular stoichiometry and longitudinal organization of meiotic chromosome axes and loops. Plos Biol 18, e3000817.

Wongso, D., Dong, J., Ueda, H., & Kitaguchi, T. (2017). Flashbody: A Next Generation Fluobody with Fluorescence Intensity Enhanced by Antigen Binding. Anal Chem 89, 6719–6725.

Wutz, G., Várnai, C., Nagasaka, K., et al. (2017). Topologically associating domains and chromatin loops depend on cohesin and are regulated by CTCF, WAPL, and PDS5 proteins. Embo J 36, 3573–3599.

Yang, X., Boehm, J.S., Yang, X., et al. (2011). A public genome-scale lentiviral expression library of human ORFs. Nat Methods 8, 659–661.

Zacharias, D.A., Violin, J.D., Newton, A.C., & Tsien, R.Y. (2002). Partitioning of Lipid-Modified Monomeric GFPs into Membrane Microdomains of Live Cells. Science 296, 913–916.

Zhao, N., Kamijo, K., Fox, P.D., Oda, H., Morisaki, T., Sato, Y., Kimura, H., & Stasevich, T.J. (2019). A genetically encoded probe for imaging nascent and mature HA-tagged proteins in vivo. Nat Commun 10, 2947.

